# Changes in oscillations in anterior cingulate and medial prefrontal cortex are associated with altered signatures of Bayesian predictive coding in trait anxiety

**DOI:** 10.1101/2022.05.24.493280

**Authors:** Thomas P Hein, Zheng Gong, Marina Ivanova, Tommaso Fedele, Vadim Nikulin, Maria Herrojo Ruiz

## Abstract

Recent advances in the computational understanding of decision-making processes have led to proposals that anxiety biases how individuals form beliefs and estimate uncertainty. The anxiety and decision-making circuitry broadly overlap in regions such as the medial prefrontal cortex (mPFC), anterior cingulate cortex (ACC), and orbitofrontal cortex (OFC). Changes in activity across these brain areas could help explain how misestimation of uncertainty and altered belief updating can lead to impaired learning in anxiety. To test this prediction, this study built on recent progress in rhythm-based formulations of Bayesian predictive coding to identify sources of oscillatory modulations across the ACC, mPFC, and OFC that are associated with altered learning in subclinical trait anxiety. In a magnetoencephalography (MEG) experiment, two groups of human participants pre-screened with high and low trait anxiety (HTA, LTA: 39) performed a volatile probabilistic reward-based learning task. We modelled behaviour using a hierarchical Bayesian learning model. Furthermore, we quantified the parametric effects of trial-wise estimates of unsigned precision-weighted prediction errors (pwPEs) and, separately, precision weights and surprise on source-reconstructed MEG time-frequency responses using convolution modelling. We showed that HTA interferes with overall reward-based learning performance associated with more stochastic decisions and more pronounced lose-shift tendencies. These behavioural effects were explained by an overestimation of volatility and faster belief updating in HTA when compared to LTA. On a neural level, we observed enhanced gamma responses and decreased alpha/beta activity in HTA during the encoding of unsigned pwPEs about about stimulus outcomes relative to LTA. These effects emerged primarily in the ACC and dorsomedial PFC (dmPFC), and they were accompanied by additional ACC alpha/beta modulations representing differential encoding of precision weights in each anxiety group. Our study supports the association between subclinical trait anxiety and faster updating of beliefs in a volatile environment through gamma and alpha/beta activity changes in the ACC and dmPFC.

## Introduction

Anxiety is a psychological, physiological, and behavioural state characterised by worry about undetermined events with potentially adverse outcomes (Grupe and Nitschke 2013; Tovote et al. 2015; Carleton 2016). A central feature in clinical and subclinical anxiety is difficulty dealing with uncertainty, playing a role in diagnosis and treatment (Quintana et al. 2016; Carleton et al. 2012; Boswell et al. 2013; Gentes and Ruscio 2011) as well as in the modelling of anxious responses (Grillon et al. 2019; Grupe and Nitschke 2011; Aylward et al. 2019). Computational modelling work has revealed that anxiety impairs learning and decision making when the associations between responses and their outcomes change due to environmental instability (environmental uncertainty or volatility; Browning et al. 2015; Pulcu and Browning 2017; Huang et al. 2017; Hein et al. 2021). Misestimation of other forms of uncertainty can also account for attenuated learning in anxiety, as shown in temporary anxiety states and in the somatic (“physiological”) component of trait anxiety (Wise and Dolan, 2020; Hein et al., 2021; Fan et al., 2021). These empirical findings converge with proposals that associate affective disorders with misestimation of uncertainty (Pulcu & Browning, 2019). Yet evidence on the neural processes associated with these computational alterations in anxiety is scarce.

Here, we build on recent progress in rhythm-based formulations of Bayesian predictive coding (PC) to identify sources of oscillatory modulations associated with altered learning in a volatile environment in subclinical trait anxiety. In a Bayesian PC framework, belief updates are informed by the discrepancy between predictions and outcomes—represented as prediction errors (PEs)—and weighted by precision (inverse variance or uncertainty of a belief distribution; Rao and Ballard 1999; Friston 2005; Bastos et al. 2012). The normative hierarchical updating policy of PC is thought orchestrated by distinct neural frequencies at particular cortical layers (Bastos et al. 2012; Sedley et al. 2016). Evidence from human MEG/EEG and monkey local field potential (LFP) studies suggest that feedforward PE signals are encoded by faster gamma oscillations (>30 Hz), while backward descending predictions are expressed in lower alpha (8– 12 Hz) and beta-band (13–30 Hz) oscillations (Bastos et al. 2012; Van Kerkoerle et al. 2014; Arnal and Giraud 2012; Auksztulewicz and Friston 2016; Bauer et al. 2006; Bastos et al. 2020). Animal studies have shown this asymmetry between alpha/beta-band power in infragranular layers and gamma-band power in supragranular layers, with alpha/beta functionally inhibiting the processing of sensory input spiking, suppressing gamma oscillations (Xing et al. 2012; Roberts et al. 2013; Bastos et al., 2015; Michalareas et al. 2016). Additional work has revealed that precision is also encoded in alpha and beta oscillations (Palmer et al. 2019; Sedley et al. 2016). As precision values weight the transmission of PEs (Feldman and Friston 2010), the composite precision-weighted PE (pwPE) signal may, as recent work suggests, be represented in antithetical modulation of gamma and alpha/beta power (Auksztulewicz et al. 2017).

Although the oscillatory correlates of PC have been primarily investigated in the sensory domain, recent evidence demonstrates that a similar mechanism in the medial prefrontal cortex (mPFC) can explain decision-making processes during exploration-exploitation (Domenech et al., 2020). In reward-based learning tasks, we recently found that beta oscillations were atypically increased in state anxiety during the encoding of revelant pwPE signals (Sporn et al. 2020; Hein and Ruiz 2022). In Hein and Ruiz (2022) there was also preliminary evidence for amplified beta activity maintaining (biased) predictions about the tendency of a stimulus-reward mapping in state anxiety. The role of gamma oscillations in mediating altered learning in anxiety through prediction error signalling remains, however, speculative. Due to the antithetic nature of gamma and alpha/beta activity in the human and non-human primate cortex (Bastos et al., 2018, 2020; Schmidt et al., 2019; Lundqvist et al., 2020), it can be predicted that anxiety-related changes in alpha and beta activity during encoding pwPE should be accompanied by opposite effects in gamma. Moreover, given the relevance of precision weighting signals (decoupled from PE values) in explaining a manifold of psychiatric conditions (Lawson et al. 2014; Adams et al. 2013; Williams et al., 2016; Friston et al. 2017; Haarsma et al. 2021), we expect that diminished or amplified precision weighting in anxiety during learning will be associated with changes in 8-30 Hz activity. This would result in biased predictions in this condition, possibly reflected in changes in alpha and beta oscillations.

The contribution of different brain regions to the frequency-domain expression of computational learning alterations in anxiety remains largely unknown. We hypothesise that neural sources that overlap with the neural circuitry of anxiety, decision making under uncertainty and reward-based learning, including the ventromedial, dorsomedial PFC (vmPFC, dmPFC), orbitofrontal cortex (OFC), and anterior cingulate cortex (ACC), will play a crucial role in the expression of altered oscillatory correlates of Bayesian PC during decision making in anxiety (Paulus et al. 2004, Grupe and Nitschke 2013; Hayden et al., 2011; Hunt et al., 2018; Robinson et al., 2019; Rouault et al. 2019; Domenech et al., 2020, Rolls et al., 2022).

Here we test these hypotheses using computational modelling and source-level analysis of oscillatory responses in MEG. We investigated a low and high trait anxious group on a binary probabilistic reward-based learning task under volatility. To assess whether trait anxiety interferes with reward-based learning performance through biased estimates of different forms of uncertainty, we modelled behavioural responses using a validated hierarchical Bayesian model, the Hierarchical Gaussian Filter (HGF; Mathys et al. 2011; Mathys et al. 2014). We then extracted HGF estimates of unsigned pwPEs about stimulus outcomes, representing precision-weighted surprise about new information, and separately, the precision ratio with which the PE is weighted. These trajectories were used as input to two separate convolution models to estimate the time-frequency responses modulated by these computational learning quantities (Litvak et al. 2013). The convolution models were solved in the reconstructed source space using beamforming (van Veen et al. 2004). This allowed us to identify the rhythm-based signatures of altered Bayesian PC in anxiety and to determine whether these emerge in brain regions previously associated with clinical and subclinical anxiety, reward-based learning and decision making (Paulus et al. 2004, Grupe and Nitschke 2013; Robinson et al., 2019; Rouault et al. 2019, Domenech et al., 2020).

## Results

### Initial learning adaptation in trait anxiety

Thirty-nine participants (24 female, 15 male) completed a probabilistic binary reward-based learning task in a volatile learning setting (Behrens et al. 2007; de Berker et al. 2016; Iglesias et al. 2013), while we recorded their neural activity with MEG. Similarly to Hein et al. (2021), participants had to learn the probability that a blue or orange image in a given trial was rewarding (outcome win, 5 points reward – outcome lose, 0 points; complementary probabilities for both stimuli, p, 1-p; **Figure 1A**). Participants were informed that the total sum of all their points would translate into a monetary reward at the end of the experiment. The task consisted of two task blocks with a total of 320 trials (TB1, 160 trials; TB2, 160 trials). The stimulus-outcome contingency mapping changed 10 times across the total 320 trials every 26 to 38 trials, with each contingency occurring twice, once in each block. The possible contingencies were 0.9/0.1 and 0.1/0.9, 0.7/0.3 and 0.3/0.7, and 0.5/0.5 (**Figure 1B)**, as in Hein et al. (2021) and de Berker et al. (2016).

**Figure 1.**
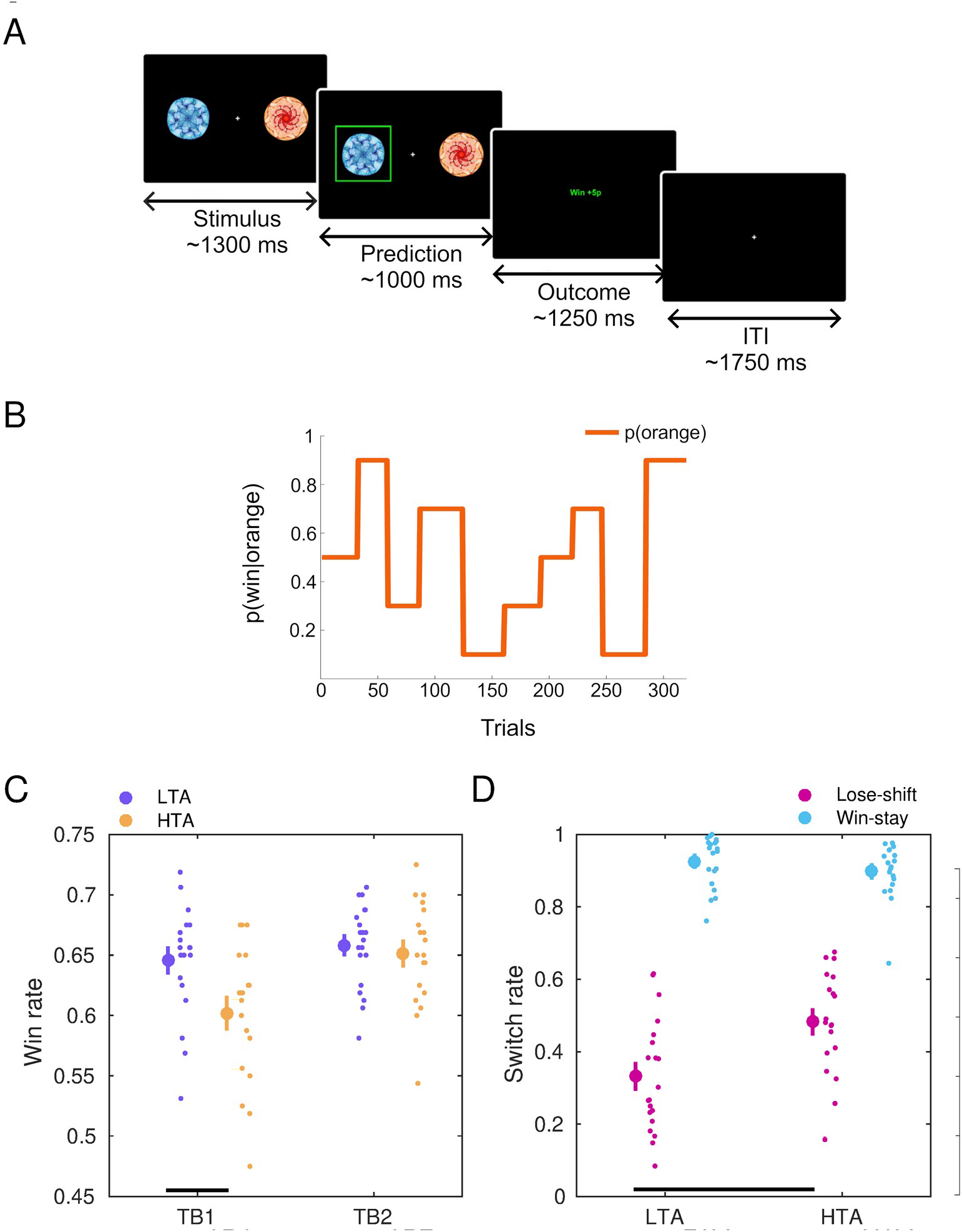
Trait anxiety modulates the win rate and the win-stay/lose-shift rates during reward-based learning. **A)** Behavioural task structure. Participants were instructed to predict which of two images was the rewarding stimulus (win = 5p) on the current trial. The stimuli (blue or orange fractal) were randomly presented to either the left or right of the screen. They remained on the screen until a response was provided or the trial *timed out* (1300 ms ± 200 ms)— recorded as no-response. After they provided a left or right side response, they immediately saw their chosen image highlighted in bright green, which remained on screen for 1000 ms (±200 ms) before the outcome was displayed. The outcome, either win or lose, was shown in the middle of the screen for 1250 ms (±200 ms) in green and red respectively. Each trial ended with a fixation cross and an inter-trial interval (ITI) of 1750 ms. **B)** The probability governing the likelihood of the orange stimulus being rewarded (p(win|orange) in one participant. **C)** High trait anxiety (HTA, yellow; N = 19) and low trait anxiety (LTA, purple, N = 20) modulated win rates differently as a function of the Block factor (significant interaction, *P* = 0.0114, 2 x 2 factorial analysis with synchronised rearrangements). There were also significant main effects of Block and Group. Post-hoc analyses demonstrated a significantly lower win rate in HTA relative to LTA in the first task block (TB1, *P_FDR_* < 0.05, permutation tests; denoted by the black bar at the bottom), but not in TB2. Within-group analyses further revealed that HTA participants significantly improved their win rate from block 1 to 2 (*P_FDR_* < 0.05, paired permutation test), whereas the win rate did not change significantly in LTA (*P_FDR_* > 0.05). Data in each group are represented using the average (large dot) with SEM bars. To the right are individual data points to display dispersion. **D)** Win-stay (blue) and lose-shift (red) rates in each anxiety group. The rates were estimated as the number of trials in that category relative to the total number of trials in the outcome type (e.g. lose-shift rate: number of lose-shift trials divided by the total number of lose events). High trait anxiety was associated with greater lose-shift rates relative to LTA (*P_FDR_* < 0.05, permutation tests), while win-stay rates did not significantly change as a function of anxiety (*P_FDR_* > 0.05). Between-group differences are marked by the bottom (lose-shift) bars.

To assess our hypotheses that trait anxiety modulates belief updating during decision making in a volatile environment, we pre-screened the participants to form two experimental groups: low trait anxiety (LTA, which we defined as score below 35 in the trait subscale of the Spielberger State Trait Anxiety Inventory, STAI, range 0-80; Spielberger, 1983; see **Materials and Methods**) and high trait anxiety (HTA, defined as a STAI trait score above 45). STAI trait scores above 45 have been associated with clinically significant anxiety in earlier work (Fisher and Durham, 1999). This supports that our HTA group corresponded with previously reported high levels of anxiety. Both LTA and HTA samples were matched in age and the proportion of males and females. During performance, our participants were monitored for physiological changes in heart-rate variability (HRV and high-frequency HRV [HF-HRV]), to control for potential confounding factors that could modulate task completion (See **Materials and Methods: Measures of anxiety**). This analysis confirmed that there were no alterations in physiological responses previously associated with temporary states of anxiety (**Supplementary file 1; Figure 1 – figure supplement 1;** Friedman, 2007; Fuller, 1992; Klein et al., 1995; Miu et al., 2009; Pittig et al., 2013). Physiological responses did not vary as a function of the task block either (**Supplementary file 1; Figure 1 – figure supplement 1).** Accordingly, most of our standard behavioural and computational measures were assessed between groups, after collapsing the block information (see **Statistics**). However, because in our recent work we observed a large effect of the task block on behavioural win rates in anxiety (Hein et al., 2020), we assessed this variable as a function of the Group (LTA, HTA) and Block (TB1, TB2) factors (**Statistics**).

Participants in each anxiety group exhibited different win rates (percentage of rewarded trials) depending on the task block (significant interaction effect of Block and Anxiety, *P* = 0.0114; non-parametric 2 x 2 factorial test with synchronised rearrangements, 5000 permutations; Basso et al., 2007). In addition, we observed a significant main effect of Block (*P* = 0.0036), and a significant Group effect (*P* = 0.0280, **Figure 1C**). Follow-up post-hoc analysis with pair-wise permutation tests revealed a significantly smaller win rate in HTA during TB1 relative to LTA (*P =* 0.015, significant after control of the false discovery rate across multiple post-hoc tests, hereafter denoted by *P*_FDR_ < 0.05; non-parametric effect size estimator, Δ = 0.73, CI = [0.64, 0.89], see **Materials and Methods**). By contrast, during the second block there was no significant between-group difference (*P*_FDR_ > 0.05, **Figure 1C**). In addition, HTA individuals exhibited a pronounced increase of the win rate from block 1 to 2 (*P*_FDR_ = 0.0036 < 0.05, paired permutation test; paired-samples effect size Δ_sup_ = 0.74, CI = [0.65, 0.87]), while this effect was not observed in LTA (*P*_FDR_ > 0.05). Crucially, the individual and group average win rates were well below the ceiling win rate (mean 0.74 [SEM 0.001], maximum 0.76, measured from the true reward contingency settings). These results demonstrate that HTA exhibited poorer reward-based learning performance relative to LTA mainly due to differences in block 1, suggesting an initial adaptation deficit. HTA individuals, however, improved considerably during block 2 leading to higher win rates that failed to differ significantly from rates in LTA.

High win rates in a fast-changing environment could be associated with a tendency to express win-stay/lose-shift behaviour more (Huang et al. 2017; Jiang et al. 2018; Xia et al. 2021). To assess this, we calculated the win-stay and lose-shift rates, which were normalised separately for each outcome type: win or lose (e.g. Grogan et al., 2007). In HTA we found a significantly higher lose-shift rate when compared with LTA (*P_FDR_* = 0.0034 < 0.05, Δ = 0.76, CI = [0.58, 0.89], **Figure 1D**), but no significant differences in the win-stay rate (*P_FDR_* > 0.05). The higher lose-shift rate in HTA relative to LTA was strikingly similar across contingency phases (90/10-10/90, 70/30-30-70 and 50/50 mappings; **Figure 1 – figure supplement 2**). Thus, across the experiment, high trait anxiety individuals consistently switch more than LTA individuals after losing in a trial.

Examining the total switch rate (trial-to-trial response switches of any kind, Aylward et al. 2019) as a proxy metric of exploratory response choices, several authors have reported impaired switching strategies in anxiety (Ansari et al. 2008; Xia et al. 2021; Paulus et al. 2004; Huang et al. 2017; Aylward et al. 2019). We therefore additionally assessed this measure and found that the overall switch rate was significantly higher in HTA than LTA (*P_FDR_* = 0.0134 < 0.05, Δ = 0.71, CI = [0.55, 0.82]; mean switch rate in each group and SEM: 0.24 [0.02] in HTA, 0.16 [0.02] in LTA). This result was, however, mainly accounted for by group differences in lose-shift rates, as shown above.

Control analyses demonstrated that the effects of trait anxiety on win rates during performance were not accompanied by changes in reaction time (RT) or by classical attentional effects that could lead to a differential slowing of RT in predictable (0.9-0.1) and unpredictable (0.5-0.5) contingency mapping phases (Prinzmetal et al., 2009; see **Supplementary file 2**). In addition, based on the Group x Block interaction effect observed in win rates, we conducted a post-hoc analysis to explore whether the lose-shift rates or total switch rates could also be explained by a Group x Block interaction. This was not the case: no significant effect of Block or Interaction effect was found in either case (*P* > 0.05), but the significant Group effect on lose-shift and total switch rates remained (*P_FDR_* = 0.0034 and 0.0120, respectively). This demonstrates that the increased tendency to shift following lose outcomes in HTA relative to LTA, like for the general switch tendency, did not change throughout the experiment, despite HTA exhibiting an initial adaptation deficit (expressed in lower win rates in TB1) that was overcome towards second block.

### Differential effects of trait anxiety on learning are best described by a hierarchical Bayesian model wherein decisions are driven by volatility estimates

We next aimed to determine whether learning differences in our anxiety groups could be accounted for by changes in estimates of different forms of uncertainty. Overestimating uncertainty in the environment may lead to anxious avoidance responses and individuals missing out on invaluable safety signals and rewarding feedback (Aylward et al. 2019; Pulcu and Browning 2019; Bublatzky et al. 2017; Maner et al. 2007). Alternatively, higher levels of estimated environmental uncertainty may inflate the degree to which new outcomes update beliefs (Lawson et al. 2017; Jepma et al. 2016). Learning can also be influenced by a different form of uncertainty, related to our imperfect knowledge about the true states in the environment (informational uncertainty; Mathys et al., 2014).

To assess different forms of uncertainty in our task, we modelled decision-making behaviour with the Hierarchical Gaussian Filter (HGF, Mathys et al. 2011; Mathys et al. 2014). This model describes hierarchically structured learning across various levels (*1,2,…, n*) and and trials *k*, corresponding to hidden states of the environment *x_1_^(k)^, x_2_^(k)^,…, x_n_^(k)^* and defined as coupled Gaussian random walks (**Figure 2A**). On level 2, *x_2_^(k)^* denotes the current true probabilistic mapping between stimulus and outcome. In our modelling approach an agent would also infer the rate of change of the tendency towards a contingency mapping, that is, the level of environmental volatility on trial *k*. This is represented by the hidden state *x_3_^(k)^*. In the HGF, higher levels of volatility would be associated with faster learning about level 2, whereas a stable environment would attenuate learning about the reward contingencies. In the following we drop the trial index *k* for simplicity.

**Figure 2.**
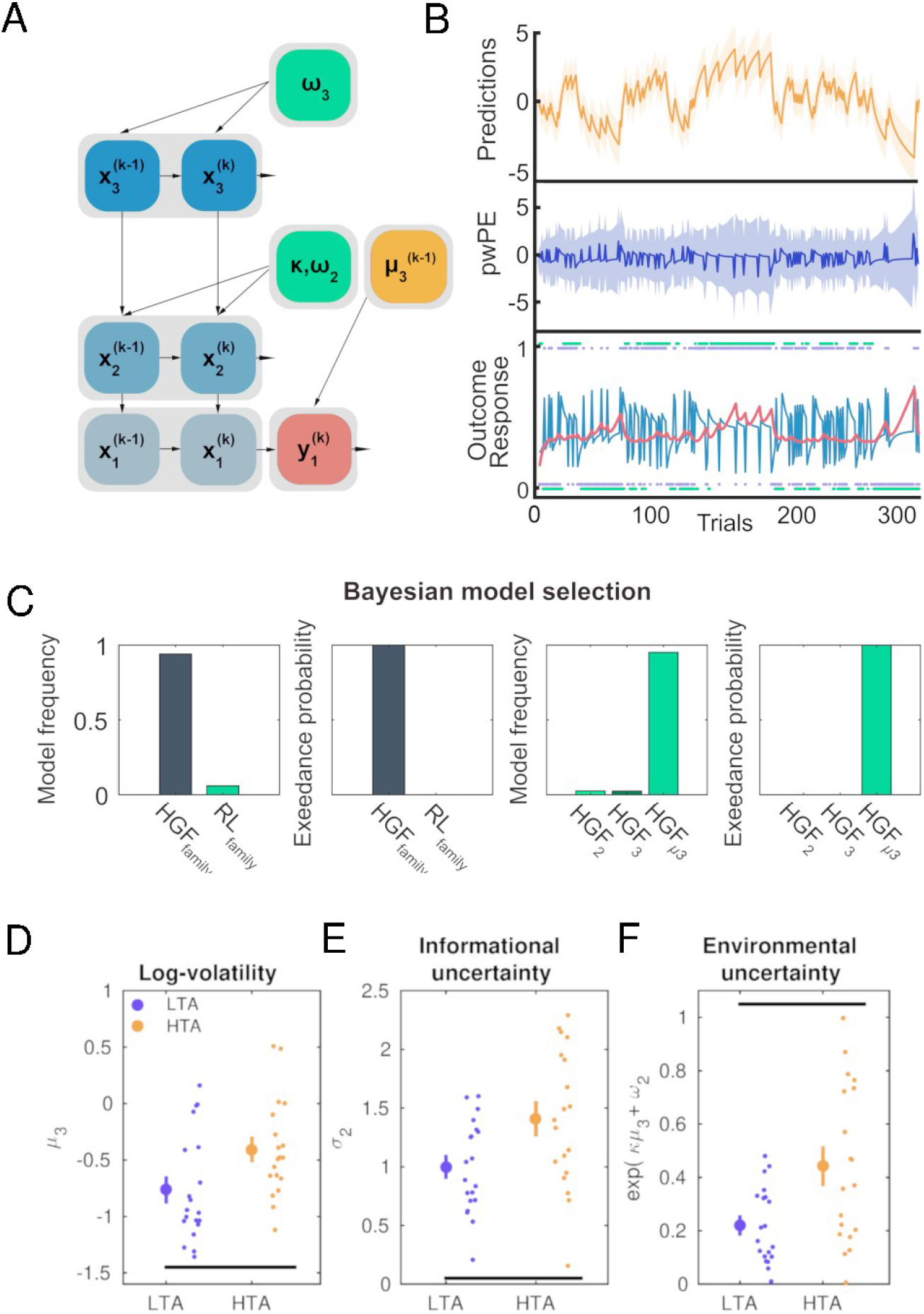
Hierarchical Gaussian Filter: Winning model and results. **A)** Representation of the three-level Hierarchical Gaussian Filter for binary outcomes with decision noise being a function of the log-volatility estimate μ_3_^(k-1)^. **B)** Associated trajectories of relevant HGF outputs across the total 320 trials in a representative participant. At the lowest level, the inputs u correspond to the rewarded outcome of each trial (1 = blue, 0 = orange; shown as purple dots). The participant’s responses y are shown in green dots tracking those trial outcomes. The light blue line indicates the series of prediction errors (PE) about the stimulus outcome, and the salmon pink line the precision weight on level 2. The middle layer of B) shows the trial-wise HGF estimate of pwPE about stimulus outcomes (pwPE updating level 2, simply termed pwPE in the graphic, ε_2_ in the main text; deep blue). For our first GLM convolution analysis, we used unsigned values of ε_2_ as the parametric regressor. The precision ratio included in the ε_2_ term weights the influence of prediction errors about stimulus outcomes on the expectation of beliefs on level 2. Predictions about the tendency towards a stimulus-reward contingency on level 2 (*μ̂*2) are displayed on the top level (yellow). We took the absolute values of this quantity as our parametric regressor (labeled simply Predictions in the graphic) in a separate exploratory GLM analysis. **C)** Bayesian model selection (BMS). The panels to the left show the model frequency and exceedance probability for the family of models ‘HGF Fam’ consisting of the 2-level HGF (HGF_2_), the 3-level HGF (HGF_3_), and the 3-level HGF informed by trial-wise estimates of volatility (HGF_μ3_) given in dark blue. The family of reinforcement learning models ‘RL FAM’ (Rescorla-Wagner, RW, Sutton K1, SK1) is presented in green. The family of HGF models better describes task learning behaviour. The right-hand panels display the model frequency and exceedance probability obtained in BMS for the three HGF models (HGF_2_: light green, HGF_3_: dark green and HGFμ_3_: green). The HGFμ_3_ model best explained the data. **D)** Log-volatility estimates (μ_3_) in HTA (yellow) were significantly higher relative to LTA (dark blue, *P*_FDR_ = 0.019 < 0.05, denoted by the black line at the bottom). **E)** Informational (estimation) belief uncertainty about the stimulus outcome tendency was greater in HTA compared with LTA (*P*_FDR_ = 0.0138 < 0.05; shown using the black bar on the x-axis). **F)** The HTA individuals were also significantly more uncertain about the environment (*P*_FDR_ = 0.0052 < 0.05, as given by the bottom black bar). No significant differences were found in uncertainty about volatility (σ_3_) or the tonic learning rates at levels 2 (ω_2_) and 3 (ω_3_).

Variational Bayesian inversion of the model provides the trial-wise trajectories of the beliefs, which correspond to the posterior distribution of beliefs about *x_i_* (i = 2,3) and represented by their sufficient statistics: *μ_i_* (mean, commensurate to a participant’s expectation) and *σ_i_* (variance, termed informational or estimation uncertainty for level 2; uncertainty about volatility for level 3; inverse of precision, *π_i_* see **Figure 2B**). On the third level, *μ_3_* represents the posterior expectation on log-volatility *x_3_*. Belief updating on each level is driven by PEs modulated by precision ratios, weighting the influence of precision or uncertainty in the current level and the level below. Formally, the update equations of the posterior estimates for level *i (i* = 2 and 3) take the following form:

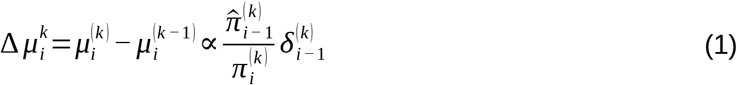

This expression illustrates that the expectation of the posterior mean on level *i*, *μ_i_^(k-1)^*, is updated to its current level *μ_i_^(k)^* proportionally to the prediction error of the level below, δ*_i-1_^(k)^*. The influence of the PE on the belief updating process is weighted by the precision ratio: the precision of the prediction of the level below 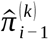, before seeing the input; divided by the precision of the posterior expectation on the current trial and level 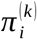). In the HGF for binary outcomes, the precision ratio updating beliefs on level two in equation (1) is reduced to *σ_2_^(k)^*. Accordingly, the posterior mean of the belief about the stimulus-reward contingencies is updated via PE about stimulus outcomes, and scaled by the degree of informational uncertainty.

To assess which model explained the trial-by-trial behavioural data in our participants better, we considered four learning models (typically labeled “perceptual models”): Two HGF models and two reinforcement learning models (Hein et al., 2021). One of the HGF models was combined with two alternative forms of response models, describing different ways in which participants’ beliefs are mapped to decisions. This resulted in a total of five alternative perceptual+response models. First, we used a 3-level HGF for binary outcomes (Mathys et al., 2014). This model describes learning about the tendency towards reward for blue/orange stimuli (level 2) and the rate of change in that tendency (volatility; level 3). Here the mapping from beliefs to responses was described by the unit-square sigmoid function and governed by a parameter ζ that regulates the steepness of the sigmoid, and is interpreted as inverse decision noise (Iglesias et al., 2013; Mathys et al., 2014). Next we used the same 3-level HGF as perceptual model with a different response model where decisions are informed by trial-wise estimates of volatility (termed HGF*_μ_*_3_; Diaconescu et al., 2014). In addition, we used a 2-level reduced HGF with a fixed volatility level (HGF_2_). We complemented our analysis by including two widely used reinforcement models, a Rescorla Wagner (RW, Rescorla and Wagner, 1972) and Sutton K1 model (SK1, Sutton, 1992). Details on the fixed and estimated model parameters are provided in **Materials and Methods,** and the prior settings are listed in **Table S1**.

We evaluated the model space using random effects Bayesian model selection (BMS, **Materials and Methods**). BMS was conducted first on the family level (HGF versus reinforcement-learning models), using the individual log-model evidence (LME) values to estimate the log-family evidence (LFE). The family of Bayesian models yielded stronger evidence (exceedance probability = 1; expected frequency = 0.94, **Figure 2C**) for explaining the task behaviour when compared with the reinforcement-learning models. Afterwards, we used the individual LME values to assess each Bayesian model (HGF_3_, HGF_2_, and HGF*_μ_*_3_). This procedure demonstrated that the HGF*_μ_*_3_ model was more likely to explain the behavioural data among participants (exceedance probability = 1; expected frequency = 0.95, **Figure 2C**). Importantly, we confirmed that the HGF*_μ_*_3_ model was equivalently the best model at describing responses independently in each group (LTA exceedance probability = 1; expected frequency = 0.91; HTA exceedance probability = 1; expected frequency = 0.91).

In the winning HGF*_μ_*_3_ response model a greater expectation on log-volatility for the current trial (defined as the estimate on trial *k-1*) is associated with a lower inverse decision temperature parameter 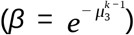, leading to a noisier mapping between beliefs and responses. On the other hand, when a participant has a lower expectation on volatility governing the stimulus-reward contingencies, she will exhibit a more deterministic coupling between her current belief and subsequent response (Diaconescu 2014). In the context of trait anxiety, the BMS result demonstrates that inferring the underlying environmental statistics and deciding upon responses is best described by a hierarchical model in which the mapping from beliefs to responses is a function of the prediction of volatility.

### Overestimation of environmental volatility in high trait anxiety

We observed higher mean log-volatility estimates, μ_3_, in the HTA group relative to the LTA group (*P*_FDR_ = 0.019 < 0.05, Δ = 0.74, CI = [0.55, 0.87]; **Figure 2D**). The higher posterior mean on log-volatility in the high trait anxiety group represents an estimated greater level of task environmental change in comparison to the low trait anxiety group. No difference was found between groups in the associated third-level parameter ω_3_ (*P* > 0.05). The result of a higher volatility estimate in HTA relative to the LTA group suggests that choice probability in HTA individuals is more stochastic. In other words, compared to LTA, HTA participants chose more often responses that were less likely to be rewarded based on their predictions for the trial.

The increased response stochasticity in HTA converges with our findings on lose-shift rates, which demonstrated an overall higher tendency to switch in HTA following lose trials—even if this goes against the current belief on the tendency of the stimulus-reward contingency. It is also aligned with the related finding of a higher overall switch rate (how often a participant changes a response independently of the outcome) in high trait anxiety individuals. As a post-hoc analysis, we conducted a non-parametric correlation across all participants between the overall switch rate and the average estimate of log-volatility, μ_3_. We found a significant association between both variables, as expected (non-parametric Spearman rank correlation ρ = 0.89, *P* < 0.00001; N = 39). Our behavioural findings thus concur with the modelling results showing that HTA individuals exhibit an overestimation of volatility, which in the HGF*_μ3_* leads to more ‘stochastic’ switching responses, which is mainly driven by switching following a lose outcome.

### Misestimation of different types of uncertainty in trait anxiety can promote learning despite an initial adaptation deficit

Informational uncertainty about the stimulus-outcome contingency, σ_2_, drives the pwPEs updating level 2, with larger σ_2_ values contributing to greater update steps. Participants with high trait anxiety had greater informational uncertainty than LTA individuals (*P*_FDR_ = 0.0138 < 0.05, Δ = 0.72, CI = [0.55, 0.86], **Figure 2E**). The overestimation of informational uncertainty in the HTA group suggests that new information has a greater impact on the update of beliefs about the tendency towards a stimulus-reward contingency (level 2), promoting faster learning on that level.

An additional important type of uncertainty governing learning in our task is uncertainty about the task environment, termed environmental uncertainty: exp(κμ_3_^(k-1)^ + ω_2_). This type of uncertainty is a function of the tonic volatility, ω_2_, and the trial-by-trial mean expectation on log-volatility μ_3_^(k-1)^ (Mathys et al. 2014). Similarly to the results on estimation uncertainty, σ_2_, we found that the HTA group had greater environmental uncertainty when compared with LTA participants (*P*_FDR_ = 0.0052 < 0.05, Δ = 0.74, CI = [0.55, 0.88], **Figure 2F**). There was, however, no significant difference between groups in the related parameter ω_2_, the expected tonic uncertainty on level 2, or in uncertainty about the volatility estimate, σ_3_ (*P* > 0.05 in both cases).

### Source analysis results

Having established that high trait anxiety is associated with a relative faster update of beliefs about the tendency of the stimulus-outcome contingency through enhanced informational uncertainty and more stochastic behaviour due to higher expectation on volatility, we next aimed to identify the source-level neural oscillatory processes accompanying these computational effects. Accordingly, we assessed the neural oscillatory representations of pwPEs and precision weights during reward-based learning in our high and low trait anxiety groups. Similarly to Hein and Ruiz (2022), this was achieved using linear convolution models for oscillatory responses (Litvak et al., 2013). This approach is an adaptation of the classical general linear model (GLM) used in fMRI analysis to time-frequency (TF) data (Litvak et al., 2013), and has been successfully used in previous EEG and MEG research (Spitzer et al., 2016; Auksztulewicz et al., 2017). It allows assessing the modulation of TF responses on a trial-by-trial basis by one specific explanatory regressor while controlling for the effect of the remaining regressors included in the model. This approach is a frequency-domain version of similar approaches used in time-domain EEG analysis, such as the massive univariate deconvolution analysis approach (Ehinger and Dimigen, 2019).

Here, we used the convolution modelling approach to explain pseudo-continuous TF data (amplitude) consisting of concatenated epochs locked to the event of interest (e.g. outcome or stimulus). Following Litvak et al. (2013), the regressors were created by convolving a Fourier basis with a set of input functions representing the onsets and values of relevant events (e.g. win, lose outcome, response). The Fourier set (sine, cosine functions) is defined over a time interval of interest (peri-event time), which results in the regressors being constructed in peri-event time as well. Fitting a GLM to the pseudo-continuous TF data provides estimates of a set of regression coefficients *β*: one regression coefficient per Fourier function and frequency (defined as in the TF data) for each event. By multiplying the estimated regression coefficients with the basis functions, we can obtain a TF image that represents an impulse response function for a specific event and is defined over the same TF range as the measured TF data. This TF image has arbitrary units and can be interpreted as a deconvolved TF response to particular discrete and parametric regressors (Litvak et al. 2013).

In the main GLM, we aimed to include the HGF trajectories of pwPEs as parametric regressors (**Figure 2B**; **Materials and Methods**). As in previous work (Auksztulewicz et al., 2017; Stefanics et al., 2018), the unsigned pwPE (|ε_2_|) on level 2 was chosen as regressor, while the pwPE regressor on level 3 was excluded due to multicollinearity (**Materials and Methods**; Vanhove, 2021; Hein and Ruiz; 2022). We chose the absolute value of ε_2_ because its sign is arbitrary: the quantity x_2_ is related to the tendency of one choice (e.g. blue stimulus) to be rewarding (*x_1_* = 1); yet this choice and therefore the sign of ε_2_ on this level is arbitrary (Hein et al., 2021). The sign does not reflect lose/win outcomes. Importantly, however, the unsigned ε_2_ value reflects some degree of lose/win information, as across participants the average |ε_2_| was significantly larger in lose relative to win trials (mean |ε_2_| [SEM]: 0.78 [0.05] for lose trials, 0.30 [0.02] for win trials; *P* = 0.0002, Δ = 0.97, CI = [0.88, 0.99]).

In a second complementary GLM, we evaluated the separate effect of the precision weights (σ_2_) and unsigned PE about stimulus outcomes (|δ_1_|) on the TF responses. These quantities, combined, represent the unsigned pwPEs on level 2: |ε_2_| = σ_2_|δ_1_|. Of note, the absolute value of PEs (|δ_1_|) is often termed *surprise* (de Berker et al., 2016), as it represents the degree of surprise about new information. Similarly as ε_2_, the sign of the PEs on level 1, δ_1_, is arbitrary in the HGF for binary responses. The PE δ_1_ represents the difference between the observed outcome for an arbitrary stimulus (e.g. whether blue is rewarding) and the participant’s belief about the probability of that outcome (**Methods and Materials**). We observed that the unsigned |δ_1_| regressor reflected lose/win information to some extent: The trial-average |δ_1_| was significantly greater for lose than win trials across participants (*P* = 0.0002, Δ = 0.97, CI = [0.88, 0.99]; mean |δ_1_| [SEM]: 0.607 [0.009] for lose trials, 0.274 [0.007] for win trials).

The GLM analyses were conducted in the source space after applying linearly constrained minimum norm variance (LCMV) beamformers (Van Veen et al., 1997) to the time series of concatenated epochs of MEG data (combined planar gradiometers and magnetometers). To assess modulations in alpha (8–12 Hz) and beta (13–30 Hz) activity, the MEG data were band-pass filtered between 1–40 Hz prior to beamforming; source-level modulation of gamma activity (32-100 Hz) was evaluated using LCMV after applying a band-pass filter between 30–124 Hz.

To reduce the data dimensionality, the convolution models were estimated in a set of brain regions previously associated with anxiety, decision making and reward processing: (1) anterior cingulate cortex (ACC), (2) orbitofrontal cortex (OFC) and related ventromedial PFC (vmPFC), and (3) dorsomedial PFC (dmPFC). The ACC and medial PFC have been consistently shown to be involved in pathological and adaptive/induced anxiety, but also in emotional and reward processing and decision making (Paulus et al. 2004, Grupe and Nitschke 2013; Chavanne and Robinson, 2019; Robinson et al., 2019). Within the medial PFC, the vmPFC represents reward probability, as well as magnitude, and outcome expectations (Rouault et al., 2019; Domenech et al., 2020). The dmPFC, on the other hand, has been shown to elicit gamma activity that correlates with unsigned reward prediction errors during exploration-exploitation (Domenech et al., 2020). The OFC is also particularly relevant in our study, as it has been associated with emotional processing, reward and punishment processing (Rolls et al., 2022). In particular, the medial OFC (mOFC) encodes reward value, whereas the lateral OFC (lOFC) encodes nonreward and punishment (Rolls, Cheng, Feng 2020; Rolls et al., 2022). The vmPFC and OFC are also considered to play a central role in the “uncertainty and anticipation model of anxiety” (Grupe and Nitschke, 2013).

These regions of interest (ROIs) corresponded to five bilateral labels (10 in total) in the neuroanatomical Desikan-Killiany–Tourville atlas (DKT, Desikan et al., 2006), which we chose to parcellate each participant’s cerebral cortex using the individual T1-weighted MRI. Our ROIs corresponded to the following anatomical labels: (1) rostral and caudal ACC (**Figure 3A**); (2) lateral and medial OFC, which include the vmPFC according to some MEG studies (Yuan et al., 2021; Morey et al., 2016; Nopoulos et al., 2010; **Figure 3B)**; (3) superior frontal gyrus (SFG), representing the dmPFC (Widge et al., 2019; **Figure 3B**). In additional exploratory analyses, however, we conducted the analysis in the other labels of the DKT atlas to identify effects outside of our ROIs.

**Figure 3.**
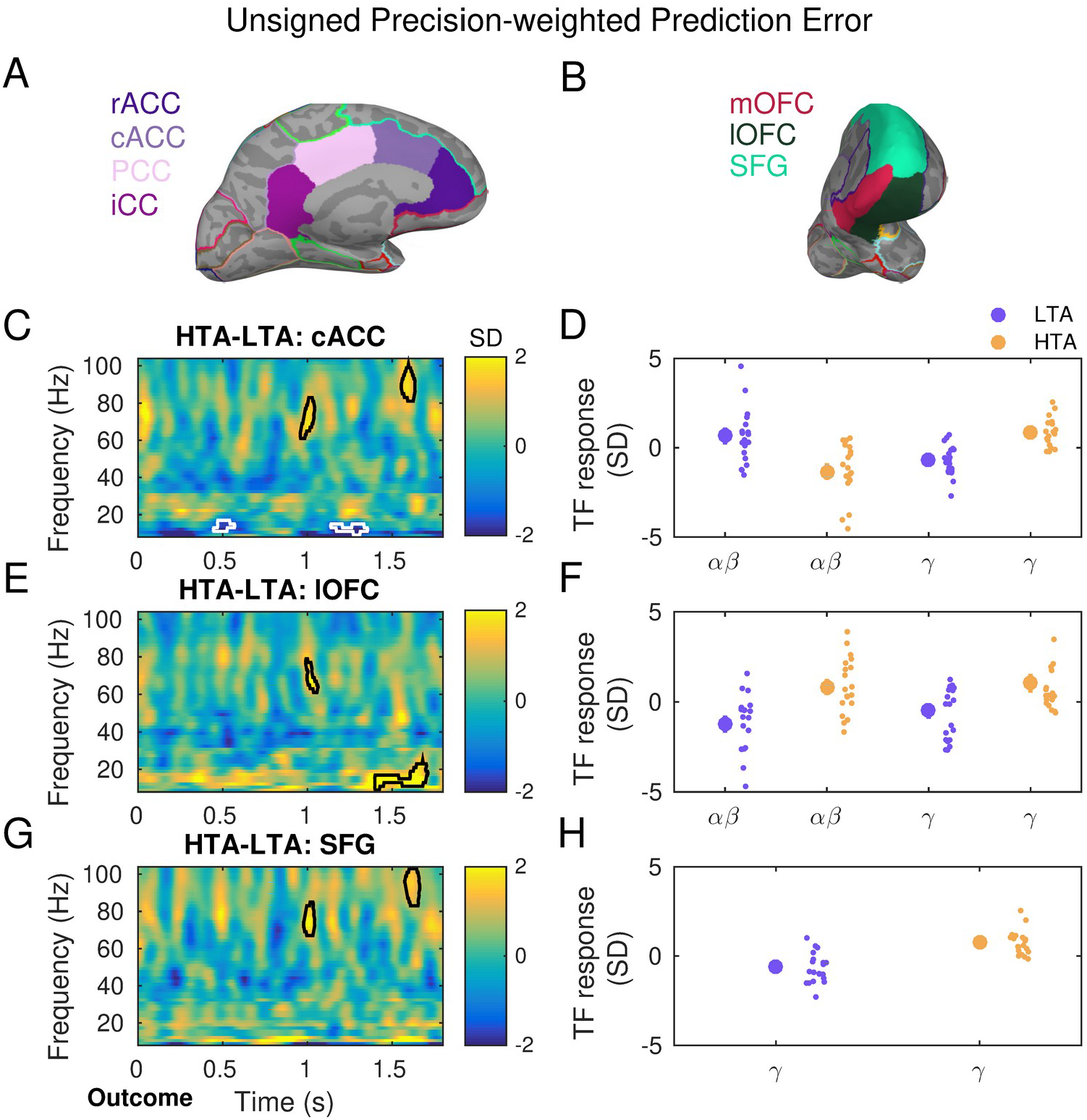
Gamma activity is modulated by unsigned precision-weighted prediction errors about stimulus outcomes and is enhanced with high trait anxiety. **A-B)** Source reconstruction of MEG signals was carried out with linearly constrained minimum norm variance (LCMV) beamforming. The statistical analysis of the convolution GLM results focused on brain regions that overlap with the circuitry of anxiety and decision making under uncertainty: ACC, OFC (lateral and medial portions: lOFC, mOFC), SFG. Panels A and B illustrate the corresponding anatomical labels in the neuroanatomical Desikan-Killiany–Tourville atlas (DKT, Desikan et al., 2006), which we chose to parcellate each participant’s cerebral cortex using the individual T1-weighted MRI. Panels **C,E,F** display between-group differences in the time-frequency (TF) images that summarise the individual oscillatory responses to the unsigned precision-weighted PEs about stimulus outcomes. TF images are shown in the 8–100 Hz range, including alpha (8–12 Hz), beta (14–30 Hz) and gamma (32–100 Hz) activity. TF images were normalised with the mean and standard deviation (SD) of the activity in a [−300, −100] ms pre-outcome interval, and thus are given in SD units. Convolution modelling and TF transformation were conducted separately in the range 8–30 Hz and 32–120 Hz in bins of 2 Hz, following LCMV beamforming. **C)** In the cACC, high relative to low trait anxiety was associated with greater gamma responses at ∼1 and 1.6 seconds during outcome feedback processing (cluster-based permutation testing, two clusters, *P* = 0.001, FWER-controlled). Significant between-group effects are denoted by the black and white contour lines in the TF images. The gamma-band effects were accompanied by a decrease in alpha-beta activity (10– 16 Hz) in HTA as compared to LTA individuals, and at 0.5 s and 1.4 s (cluster-based permutation testing, *P* = 0.01 and 0.008, FWER-controlled). **D)** The relative gamma increase in HTA shown in C) was due to more positive gamma activity during encoding unsigned pwPE in HTA than in LTA. On the other hand, HTA individuals exhibited a negative change in 10–16 Hz responses, in contrast to the positive alpha and beta activity observed in LTA individuals. This resulted in the negative between-group effect in 10–16 Hz activity in C). The large circles represent the mean (and SEM) TF response in the significant spectrotemporal clusters in C), shown separately for low frequency (alpha, beta) and gamma activity, and for each group (LTA: purple; HTA: yellow). Individual dots represent individual participant average values. **E-F)** Same as C-D) but in the lOFC. Significant clusters in gamma and 10–16 Hz (*P* = 0.01 and 0.005, FWER-controlled). **G-H)** Same as C-D) but in the SFG. Significant clusters in gamma frequencies (*P* = 0.001, FWER-controlled). rACC, rostral anterior cingulate cortex; cACC, caudal ACC; PCC, posterior CC; iCC, isthmus of the CC; SFG, superior frontal gyrus; lOFC, lateral orbitofrontal cortex; HTA, high trait anxiety; LTA, low trait anxiety.

We tested the hypothesis that high levels of trait anxiety are associated with changes in gamma and concomitant alpha/beta activity during encoding precision-weighted PE signals. In addition, we hypothesised that trait anxiety modulates alpha/beta oscillatory activity during the representation of precision weights. In Hein and Ruiz (2022), the Group effect of pwPE on neural oscillatory responses occurred approximately between 1000–1600 ms post-outcome, similarly to the latency of precision-weighting and surprise signals.

In a separate exploratory analysis we evaluated the neural oscillatory responses associated with maintaining predictions about the tendency of the stimulus-outcome contingency, *μ̂*_2_. As argued in our previous work (Hein and Ruiz, 2022), the neural representation of maintenance of predictions is likely to be diffuse in time and could be potentially observed somewhere between the stimuli presentation and the outcome resolution, co-occurring with response commission. Because the convolution approach can isolate and model overlapping events (Litvak et al., 2013), it is particularly suitable for analysing the expression of predictions in EEG/MEG data. However, identifying the neural correlates of HGF prediction trajectories is challenging as acknowledged in previous work (Diaconescu et al., 2017).

To test the hypothesis that anxiety modulates alpha/beta oscillatory activity during the maintainance of predictions, we run a separate “stimulus-locked” convolution GLM with discrete regressors denoting the stimuli presentation and the participant’s response (response: left, right, no response), and the relevant parametric regressor: the unsigned predictions on level 2 (| *μ̂*_2_|) about the tendency towards a certain stimulus-reward contingency (henceforth: ‘predictions’). As for pwPEs updating level 2, the sign of *μ̂*_2_ is arbitrary as it represents the tendency of the stimulus-reward mapping for an arbitrary stimulus (e.g. mapping for the blue image). The absolute values |*μ̂*_2_| do, however, represent a prediction about the tendency towards a particular stimulus-reward contingency. Accordingly, if a participant has a greater value of | *μ̂*_2_| in one trial, she will have a stronger expectation that given the correct stimulus choice a reward will be received. See further details in **Materials and Methods**. In Hein and Ruiz (2022), between-group differences in oscillatory responses during encoding predictions were observed 200-650 ms post-stimulus, before observing the outcome on the current trial.

### Unsigned precision-weighted prediction errors about stimulus outcomes

Our primary convolution GLM of TF responses introduced participant-specific trial-by-trial trajectories of unsigned pwPEs updating level 2, |ε_2_|, as parametric regressor, while it simultaneously controlled for the effect of discrete regressors (lose, win outcomes; no response). This GLM focused on the time interval 0–1.8 s post-outcome. A between-subject independent sample cluster-based permutation test between 8–100 Hz on the TF responses to |ε_2_| revealed a significant decrease at 10–16 Hz in the HTA group relative to LTA in the caudal portion of the ACC (two negative spectral-temporal clusters, *P* = 0.01 and 0.008, two-sided test, FWER-controlled, 3D data: 10 labels x samples x frequency bins). The latency of the significant effect was 450–550 ms and 1140–1350 ms (**Figure 3C–D**). A second significant effect in the low-frequency range was found in the lateral OFC, due to increased 10–22 Hz activity in HTA when compared to LTA participants (positive cluster within 1400–1700 s, *P* = 0.008, two-sided test, FWER-controlled; **Figure 3E–F**). Crucially, the latency of these effects extended for at least two full cycles of the central cluster frequency. In the gamma range, we observed prominent increases in TF responses in HTA as compared to LTA participants in the cACC, lOFG and SFG (positive clusters, *P* = 0.001, 0.005, and 0.001, two-sided test, FWER-controlled; **Figures 3C–H**). The enhanced gamma modulation in HTA relative to LTA had a similar latency across these regions: it emerged at around 1000 ms within 60–80 Hz and later at 1600 ms within 80–100 Hz. The gamma effects extended for at least 5 cycles at the central cluster frequency. No other effects were found.

The TF images of the discrete win and lose regressors in this outcome-locked GLM revealed similar between-group beta effects in brain regions that overlapped with those associated with the pwPE regressor (**Figure 3 – figure supplement 1**), in line with predictive coding proposals (Friston et al., 2005). Notably, however, the polarity of most of the between-group differences for the win and lose outcome regressors was reversed: we observed a relative attenuation of gamma amplitude for HTA relative to LTA in the cACC and SFG, and an additional reduction in the beta frequency range in the lOFC.

We reasoned that the greater gamma activity observed in HTA in the cACC, SFG (dmPFC) and lOFC during encoding |ε_2_| could reflect an association between larger |ε_2_| values and a greater likelihood of switching responses in HTA. In the ACC, reward and value estimates guide choices, with higher ACC activity observed in trials leading to choices (de Berker et al., 2019). In addition, activity in the dmPFC represents value difference signals modulating motor responses (Hare et al., 2011). We therefore asked whether trials leading to a response shift had larger |ε_2_| values, due faster belief updating; we also assessed whether this effect was modulated by the Group factor. A 2 x 2 Group x Shift (trial followed by a response shift / no shift) analysis of unsigned pwPE values on level 2 demonstrated a significant main effect of the Group and Shift factors (*P* = 0.0068, 0.0012 respectively; **Figure 3 – figure supplement 2).** A significant interaction effect was also observed (*P* = 0.0200). These results demonstrate that |ε_2_| was larger in trials followed by a shift in the choice made by participants; |ε_2_| was also greater in HTA participants overall. Moreover, the modulation of |ε_2_| values by the Shift factor was more pronounced in HTA. Complementing these results, the gamma response to the unsigned pwPE regressor in the cACC and dmPFC was associated with the rate of response shift across participants (non-parametric Spearman correlation: ρ = 0.6030, *P* = 0.0001 in the cACC; ρ = 0.5166, *P* = 0.0009 in the dmPFC). Accordingly, individuals with a greater gamma modulation by |ε_2_| in the cACC and dmPFC were more likely to shift their response in the following trial, even in unwarranted cases (i.e. following a lose outcome after choosing the more likely rewarded choice in a 0.9 contingency block). Gamma responses in the lOFC were not associated with the response shift rate (ρ = 0.21, *P* = 0.1396).

To control for the effect of the choice of the Fourier basis set on the high-frequency gamma modulations, we conducted a control analysis in which we increased the number of basis functions from 20 to 40 (**Materials and Methods).** This provided increased temporal resolution in convolution modelling. The statistical results were very similar (**Figure 3 – figure supplement 3**). In an additional exploratory analysis, we assessed the modulation of theta (4–7 Hz) activity, as this rhythm has been shown to be a marker of clinical and subclinical anxiety (Cavanagh and Shackman, 2015; Shadli et al., 2021). Moreover, theta activity can facilitate encoding of unpredictable stimuli (akin to PE), driving gamma activity (Bastos et al., 2020). Convolution modelling revealed a general increase in the amplitude of theta activity to the unsigned pwPE regressor in HTA when compared to LTA (**Figure 3 – figure supplement 4**; *P* < 0.05, *uncorrected*, rostral ACC and isthmus CC). The larger theta amplitude induced by the pwPE regressor was observed before and around 0.5 seconds and later at 1.5 seconds (**Figure 3 – figure supplement 4**). The lose outcome regressor, on the other hand, induced a suppression of theta activity in HTA relative to LTA participants, also in portions of the CC/ACC but additionally in the medial OFC (**Figure 3 – figure supplement 5;** *P* < 0.05, uncorrected). No other effects were observed.

### Modulation of surprise and precision weights by anxiety

The relative reduction in alpha/beta activity in HTA during encoding unsigned pwPE about stimulus outcomes (**Figure 4A**) could be associated with changes in precision weights modulating 8–30 Hz activity or, alternatively, with a modulation by unsigned PEs about stimulus outcomes—representing the *surprise* experienced by the participants (de Berker et al., 2016). In our recent work we found a convergence between the effects of |ε_2_| and surprise on 13–30 Hz beta activity in state anxiety relative to a control group (Hein and Herrojo Ruiz, 2022). Precision weights, on the other hand, were positively correlated with alpha activity (8–12 Hz) and dissociated between groups. Our second GLM thus aimed to determine the separate effect of the precision weight (σ_2_) and unsigned PE (|δ_1_|) regressors on the TF responses.

**Figure 4.**
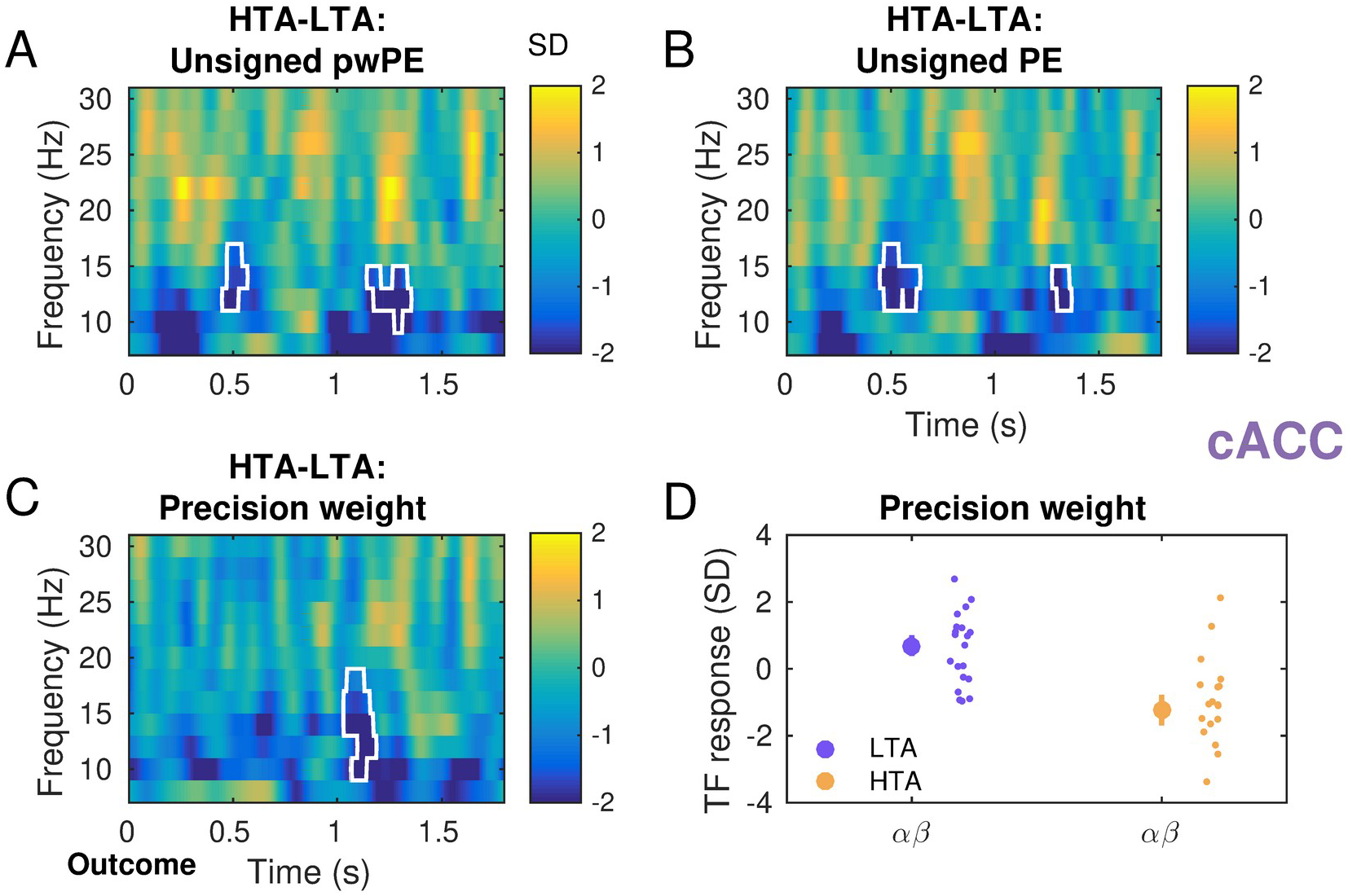
Attenuation of alpha and beta activity during the representation of precision weights and unsigned PEs in high trait anxiety. **A)** Zoom in on Figure 3C. TF images were normalised with the mean and standard deviation (SD) of the activity from −300 to −100 ms before the outcome feedback, and thus are given in SD units. In the caudal ACC (cACC) we had observed in HTA relative to LTA a reduction in 10–16 Hz activity during encoding unsigned pwPEs about stimulus outcomes (Figure 3). To investigate the effect of trait anxiety on the neural oscillatory correlates of precision weights modulating the influence that PEs have on updating predictions on level 2, we implemented a new convolution model. This GLM used as separate continuous regressors the unsigned PE about stimulus outcomes, |δ_1_| (surprise), and the precision weights on level 2, σ_2_ (informational uncertainty); the model also included discrete regressors coding for outcome events (win, lose, no response). **B)** In the cACC, the between-group effects in the TF responses to surprise were very similar to those in A). We observed a reduction in 10–16 Hz activity in HTA relative to LTA observed at 0.5 and 1.4 s (*P* = 0.01, FWER-controlled). **C)** The representation of precision weights was also associated with a relative reduction of higher alpha and low beta activity in high trait anxiety, yet at latencies that were aligned with the gamma effects in Figure 3 (1.1 and 1.6 s; *P* = 0.01, FWER-controlled). **D)** Average activity in the significant spectrotemporal clusters in C) separately for each group (LTA: purple; HTA: yellow). The large dot denotes mean and SEM as error bars. Individual dots represent individual participant average values.

Processing surprise about stimulus outcome was correlated with a reduction in 10–14 Hz activity in HTA relative to LTA in the bilateral caudal portion of the ACC (two negative clusters, *P* = 0.01, two-sided test, FWER-controlled. **Figure 4B**). The spectral and temporal distribution of this effect had a considerable overlap with the effects of the |ε_2_| regressor on 8–30 Hz (**Figure 4A**, centered at 1 s and 1.4 s, and extending for at least two full oscillation cycles). Interestingly, the TF responses to the precision-weight regressor were also attenuated in the low frequency range in high relative to low trait anxiety individuals (one negative cluster, *P* = 0.01, two-sided test, FWER-controlled. **Figure 4C**). However, the between-group effect extended up to 18 Hz in the beta band and emerged at a different latency, 1000–1200 ms, corresponding to the latency of the gamma effect of |ε_2_| (**Figure 3)** and extending for 1-2 oscillation cycles. Inspection of the individual values of 10–18 Hz activity revealed that low trait anxiety participants had predominantly a positive alpha/beta activity response to the precision-weight regressor. By contrast, high trait anxiety individuals exhibited mainly an attenuation of 10–18 Hz activity during the representation of precision weights (**Figure 4D)**.

In addition, enhanced 10–16 Hz and 12–20 Hz activity was observed in the lateral OFC in both hemispheres for the surprise and precision-weight regressors, respectively (at ∼ 1.5 s for σ_2_ and 1.6–1.7 s for surprise, |δ_1_|; *P* = 0.02 in each case, two-sided test, FWER-controlled; **Figure 4— figure supplement 1**). Exploratory analyses in anatomical labels outside of our ROIs showed a between-group effect of precision weights in alpha/beta activity exclusively in the posterior cingulate cortex (negative cluster, *P* = 0.02, two-sided test, *uncorrected*).

### Stimulus-locked predictions about reward tendency

The stimulus-locked convolution model allowed us to explore the modulation of alpha/beta oscillatory activity by the prediction regressor, while controlling for the simultaneous effect of discrete stimulus and response regressors. We observed a significant between-group difference in the beta-band TF responses to |*μ̂*_2_|, due to greater beta activity in participants with high relative to low trait anxiety (**Figure 5AB**; significant positive clusters at 100-200 ms and 600-680 ms post-stimulus, P = 0.005, FWER-controlled). This effect was limited to the caudal ACC (right and left hemisphere). This result contrasted with the pronounced drop in beta activity observed in HTA relative to LTA with the discrete stimulus regressor (**Figure 5CD;** stimuls left; similar findings for stimulus right). For the stimulus regressor, significant between-group differences were found in the beta range and extending across a series of clusters from 100 to ∼700 ms post-stimulus (*P* = 0.001, FWER-controlled; These effects emerged before the feedback presentation, which occurred at around ∼1550 ms on average across participants: 1000 [± 200] ms after the response, which had a latency of 550 [20] ms on average). All significant between-group effects extended for at least one cycle at the relevant cluster frequency.

**Figure 5.**
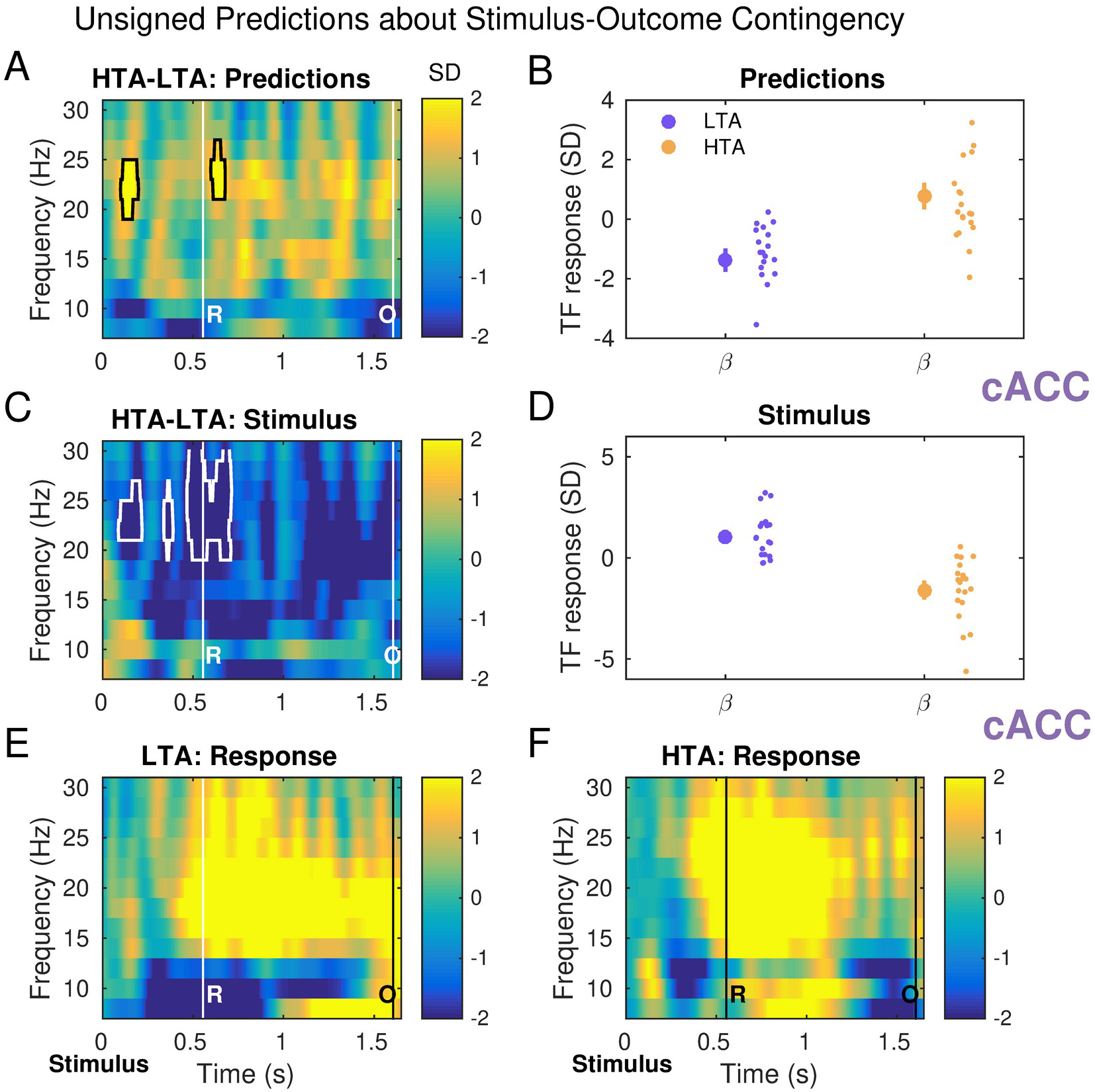
Stimulus-locked beta activity reflects differential modulation with anxiety by predictions about the reward tendency. **A)** The regressor parametrising absolute predictions about the tendency towards a stimulus-reward contingency modulated beta activity differently in HTA and LTA participants (significant positive clusters at 100-200 ms and 600-680 ms post-stimulus, *P* = 0.005, FWER-controlled). This effect was limited to the right and left caudal ACC (right cACC results shown in the figure). The average response time in the combined sample was 555 (20) ms, which is denoted by a vertical line at the corresponding latency, labeled “R”. The outcome was presented 1000 ms (± 200 ms) following the response, here denoted by the vertical line labeled “O”. **B)** Average activity in the significant spectrotemporal clusters in A) shown separately for each group (LTA: purple; HTA: yellow). The large dot denotes mean and SEM as error bars. Individual dots represent individual participant average values. **C)** In the same convolution model, we observed that the discrete stimulus regressors induced a pronounced drop in beta activity in HTA when compared to LTA in the right cACC (shown here for blue stimulus on the left; similar results for blue stimulus on the right, not shown). This reduction was maintained throughout the post-stimulus interval (significant clusters from 100 until ∼700 ms, before the feedback presentation; *P* = 0.001, FWER-controlled). **D)** Same as B) but for the stimulus (blue left) regressor. The relative reduction in beta activity shown in C) was associated with increases in the individual beta-band TF responses to the stimulus regressor in LTA participants, and reductions in HTA participants. **E-F)** Stimulus-locked analysis of the response regressor (left response). LTA and HTA groups exhibited similar TF images in association with the response regressor. Panels E and F illustrate the expected alpha reduction prior to and during the response, followed by a pronounced beta rebound effect. There were no between-group differences in the TF images to the discrete regressors in any of our ROIs or additional anatomical labels (*P* > 0.05). See **Figure 5 – figure supplement 1.**

Last, the discrete response regressors (response left, response right) induced in each group a prominent alpha reduction prior to and during the button press, and a classic subsequent beta rebound (**Figure 5EF** displays the results in the right cACC, stimulus left regressor). There were no significant between-group differences in the oscillatory responses to these regressors. Neither in our ROIs (**Figure 5 – figure supplement 1;** P > 0.05), nor in the additional DKT anatomical labels (P > 0.05).

## Discussion

Our study revealed that key brain regions of the anxiety and decision-making circuitry exhibit changes in oscillatory activity that can account for behavioural and computational effects of anxiety within a Bayesian predictive coding framework. We showed that high trait anxiety interferes with overall reward-based learning performance, which was associated with biases in different forms of uncertainty estimation. Inflated estimates of environmental volatility drove these changes, in line with previous reports that anxious learners overestimate volatility in all environments (Bishop and Gagne, 2018). Noisier decisions and more pronounced lose-shift tendencies accompanied higher volatility estimates for HTA participants.

On a neural level, convolution modelling revealed that HTA enhances gamma responses and decreases alpha/beta activity correlated with unsigned pwPEs about stimulus outcomes relative to LTA. These effects were observed in the ACC, lateral OFC, and dorsomedial PFC. The finding of increased gamma-band amplitude to unsigned pwPEs suggests that HTA individuals update their beliefs on the tendency of the stimulus-reward contingency more than LTA individuals, which promotes a response shift. As we argue below, the results on uncertainty estimates and the neural representation of precision weights support this interpretation. This study is the first to examine source-level oscillatory activity in subclinical anxiety to describe how misestimation of uncertainty can lead to impaired learning in this condition.

### Overestimation of different types of uncertainty explains more stochastic decisions and greater belief updates in trait anxiety

Current theories of affective disorders conceptualise some of the psychiatric symptoms as divergent hierarchical Bayesian inference, described by difficulties estimating uncertainty and balancing the influence of sensory input on updating prior beliefs (Pulcu and Browning 2019; Paulus and Yu 2012). These proposals extend to subclinical anxiety, given the considerable overlap of behavioural and neural effects in pathological and subclinical populations (Grupe and Nitschke, 2013; Robinson et al., 2019; Chavanne and Robinson, 2021; Shadli et al., 2022). Our results are in line with these predictions, demonstrating that individuals with HTA have a greater degree of informational uncertainty, *σ_2_*. HTA participants also had larger environmental uncertainty and overestimated environmental volatility, *μ_3_*. Greater informational uncertainty (smaller precision *π_2_*) drives faster update steps on the beliefs on the tendency of the stimulus-reward contingency (Mathys et al., 2014). Larger *μ_3_* values also influence lower-level pwPEs, inflating the degree to which new outcomes update beliefs (Lawson et al. 2017; Jepma et al. 2016). Thus, subclinical trait anxiety is associated with faster updating of beliefs about stimulus-reward contingencies through an overestimation of informational uncertainty and environmental volatility.

The computational results contrast with our recent findings on induced state anxiety, revealing an attenuation of belief updating about the reward contingencies governing the environment (Hein et al., 2021). In that work, state-anxious individuals underestimated informational and environmental uncertainty. Similar to the state anxiety results, the somatic (“physiological”) component of trait anxiety has been linked to underestimation of uncertainty and relative uncertainty between choices during exploration (Wise and Dolan, 2020; Fan et al., 2021). By having a reduced ability to tolerate uncertainty, temporarily anxious individuals or individuals high on somatic anxiety may exhibit a more limited behavioural repertoire, exploiting previous actions while missing out on potentially more rewarding choices (Sporn et al., 2021; Fan et al., 2021).

Here, biased estimates of uncertainty in HTA were associated with suboptimal switching behaviour, such as a pronounced lose-shift tendency, even in more deterministic contingency phases. A hierarchical Bayesian model in which the mapping from beliefs to responses was a function of the volatility estimate best described the participants’ behaviour in our task. As HTA participants had a larger prediction of volatility, this model implied that, compared to LTA, the HTA group chose more often responses that were less likely to be rewarded based on their predictions for the trial. Increased response stochasticity in the HTA group agrees with its larger lose-shift rate and overall switch rate. This may explain the initially poorer task performance of the HTA group, as higher levels of response switching combined with a high learning rate would make it difficult to infer the true underlying probabilistic contingencies. In this scenario, distinguishing between meaningful environmental changes and outcome randomness would be more challenging. The increased response stochasticity in high trait anxiety contrasts with the reported reduction in response exploration and choice stochasticity in state anxiety and the somatic component of trait anxiety (Sporn et al., 2021; Fan et al., 2021). By modelling unpredictability, volatility (Piray & Daw, 2020b), and subjective uncertainty (confidence ratings; as argued in Fan et al., 2021) separately, follow-up work could determine whether a subjective misattribution of the causes of loss outcomes (McDougle et al., 2015) could account for the increased choice stochasticity in anxiety.

The volatility results in our study converge with findings of Huang et al. (2017), who described an inflexible adjustment of learning rates—remaining suboptimaly large—to volatility, as well as an inflated lose-shit rate. Browning et al. (2015) similarly showed that high trait anxious individuals in an aversive learning context cannot adapt their learning rate to changes in environmental volatility. The authors interpreted these outcomes as maladaptive, concluding that HTA individuals were not sensitive to environmental change. Despite the differences in task structure and feedback valence between these studies and our own, these findings support a relative overlap in clinical and subclinical (trait) anxiety in how volatility is inferred or used to drive learning. By using a hierarchical Bayesian model that provides dynamic trial-to-trial trajectories of beliefs, our study extends previous data on maladaptive responses of anxiety to volatility, accounting for behavioural effects through changes in uncertainty estimates. Crucially, however, high trait anxiety can also result in fewer lose-shift responses during volatile probabilistic learning tasks (Paulus et al. 2004; Xia et al., 2021), yet still impair performance overall; or at least initially (Jiang et al. 2018). Current approaches to investigating transdiagnostic psychiatric symptoms in large samples of the general population (Wise et al., 2020) could be applied to study the effect of environmental volatility on learning in anxiety and explain mixed results in previous work.

### Changes in oscillatory activity in the ACC, OFC, and mPFC are associated with altered Bayesian predictive coding in anxiety

By applying convolution models to explain amplitude modulations in time-frequency MEG responses (Litvak et al., 2013, Spitzer et al., 2016), we were able to identify the effect of trait anxiety on the neural oscillatory correlates of unsigned pwPE, precision weights and surprise about stimulus outcomes. A complementary exploratory analysis produced tentative evidence on the time-frequency responses of predictions about the reward contingencies. Crucially, the convolution analysis controls for the simultaneous effect of discrete and parametric regressors as well as for variable event latencies trial-to-trial (Litvak et al., 2013). As such, this approach has been successfully used in previous work investigating neural encoding of hierarchical Bayesian quantities in the frequency domain (Auksztulewicz et al., 2017; Hein and Herrojo Ruiz, 2022). By modelling source-reconstructed time-frequency responses to HGF regressors, our work extends time-domain EEG and fMRI studies of Bayesian inference and predictive coding (Iglesias et al., 2013; Powers et al. 2017; Diaconescu et al. 2017; Stefanics et al. 2018; Nassar et al. 2019; Weber et al. 2020), providing novel insights into rhythm-based formulations of Bayesian PC (Arnal and Giraud, 2012; Bastos et al., 2015; 2020; van Pelt et al., 2016; Wang, 2010).

Encoding of unsigned pwPEs about stimulus outcomes was associated with dampened alpha/beta oscillations (10–16 Hz) in the caudal portion of the ACC in HTA relative to LTA. This effect emerged between 1100–1330 ms, converging with the latency of beta modulations during pwPE encoding in our previous studies of decision making and motor learning in state anxiety (Sporn et al., 2021; Hein and Ruiz, 2022). Temporary anxiety states, however, enhance the amplitude of beta oscillations (Sporn et al., 2021; Hein and Ruiz, 2022). An additional alpha/beta activity reduction in HTA as compared to LTA was observed earlier at 500 ms. Importantly, the alpha and beta attenuation effect was accompanied by a pronounced phasic increase in the amplitude of gamma responses in HTA and at ∼1 and 1.7 s. The relative gamma increase in the HTA group was identified across three ROIs, the cACC, the lateral OFC and the dorsomedial PFC (label SFG in the DKT atlas, Widge et al., 2019). The results are consistent with the notion that bottom-up PEs are encoded in gamma frequency oscillations (Michalareas et al. 2016; Bastos et al. 2015; Bastos et al. 2012; Arnal and Giraud 2012). In the context of trait anxiety, the results align with the computational findings on uncertainty estimates, suggesting that trait anxiety promotes outcome-driven processing, enhancing the role of PEs in updating predictions (Bauer et al., 2014; Sedley et al., 2016).

Our study is the first to demonstrate that alterations in Bayesian belief updating during reward-based learning in subclinical trait anxiety can be associated with changes in gamma activity across key brain regions of the anxiety and decision-making networks. The ACC and medial PFC have been consistently shown to be involved in pathological and induced anxiety but also decision making and the processing of rewards or emotions (Paulus et al., 2004, Grupe and Nitschke, 2013; Hunt et al., 2018; Chavanne and Robinson, 2019; Robinson et al., 2019). The gamma effects we observed had very similar latencies in the cACC and the dmPFC (SFG), whereas no effects were found in the medial OFC, which is considered to include the vmPFC in the anatomical parcellations in MEG studies (Yuan et al., 2021; Morey et al., 2016). Larger gamma activity across the cACC and dmPFC regions was associated with a higher response-shift rate across participants, and trials that were followed by a response shift had larger |ε_2_| values, more prominently in HTA. These findings are consistent with some accounts of ACC and dmPFC function, suggesting that signals in these brain regions guide response choices (Hare et al., 2011; de Berker et al., 2019). The gamma effects in the dmPFC are particularly interesting as they are aligned with recent work linking gamma oscillations in the human dmPFC to encoding unsigned reward PEs during exploration-exploitation (Domenech et al., 2020). Our results provide further evidence that the rhythm-based formulations of PC and Bayesian PC (Bastos et al., 2015; 2020; Sedley et al., 2016) can be extended beyond the perceptual domain to explain learning in more general contexts.

The antithetic modulation of alpha/beta and gamma activity by the unsigned pwPE regressor in the cACC converges with the vast evidence that increased gamma power in cortex during bottom-up processing is accompanied by a dampening of alpha/beta oscillations (Hoogenboom et al., 2006; Lundqvist et al., 2020; Lundqvist et al. 2018, 2016; Bastos et al., 2018, 2020). The lateral OFC, however, elicited a relative increase first in the gamma band and subsequently at alpha/beta frequencies (10–22 Hz). The lateral OFC plays a role in encoding punishment value, nonreward and unpleasantness (Rolls et al., 2003; 2022; Cheng et al., 2016). It is possible that the relative HTA-LTA increase in alpha/beta activity in the lOFC to the pwPE regressor is due to the simultaneous negative amplitude change at the same latency and frequency range to the lose regressor. Alternatively, the increase in alpha/beta activity in the lOFC could be excitatory, contributing to further encoding the unsigned pwPE regressor. This interpretation remains speculative, however, although there is some evidence that beta activity may be excitatory in some brain regions during encoding unpredicted inputs, as shown recently in the primate parietal cortex (region A7; Bastos et al., 2020). The pwPE results across our ROIs highlight that the anticorrelated nature of gamma and alpha/beta oscillations during encoding pwPEs are expressed in specific regions of the decision-making networks (here cACC).

Complementing the results on the neural correlates of unsigned pwPE, a second convolution model demonstrated a consistent HTA-related attenuation of alpha/beta activity during encoding precision weights in the cACC. In the HGF for binary outcomes, the precision-weight term scaling the influence of PEs on the update of beliefs about the stimulus-reward contingency is simply σ_2_, the expectation on informational uncertainty. The computational results established that high trait anxiety biases uncertainty estimates, increasing σ_2_, and decreasing the precision of posterior beliefs about the stimulus-reward contingency. The GLM results thus associate the faster belief updating in HTA with a reduction of alpha/beta activity in the cACC. This outcome could be mediated by increases in synaptic gain, as proposed for alpha oscillations in attentional tasks (Bauer et al., 2014), which would promote the transmission of PEs, in line with our gamma results. This interpretation is supported by the latency of the effects, as the alpha/beta changes with precision weights were observed at 1 and 1.6–1.7 s post-outcome, which closely matches the latency of the gamma-band effects in the cACC. The latency of 8–30 Hz modulations by the surprise regressor, on the other hand, were aligned with the timing of alpha/beta attenuation effects of pwPEs, as we observed in our recent work (Hein and Herrojo Ruiz, 2022).

Combined, the neural and computational results on precision weights provide a coherent picture of the relevance of assessing precision signaling to identify routes through which subclinical trait anxiety can hinder learning, particularly when learning is embedded in an environment rich in volatility. Our results build on the mounting evidence on the role of precision in explaining altered learning in a whole suite of clinical conditions and symptoms, such as hallucinations in Parkinson’s disease (O’Callaghan et al. 2017; Friston, 2017), schizophrenia, autism (Friston et al., 2016; Lawson et al., 2014) and psychosis (Haarsma et al., 2020). An exciting avenue of future research in anxiety would be the combination of MEG recordings with pharmacological interventions, to assess the modulatory effects of neurotransmitters (dopamine: Iglesias et al; 2013; Haarsma et al., 2020; acetylcholine: Moran et al., 2013; noradrenaline: Dayan & Yu, 2006) on the neural oscillatory correlates of precision.

## Limitations

In contrast with the pwPE and precision weighting results, which converge with the behavioural and computational findings, the exploratory analysis of predictions as a regressor revealed unexpected outcomes. Maintaining predictions about the reward contingencies was associated with larger alpha/beta activity in the cACC in HTA when compared to the LTA group. If HTA participants updated their level 2 beliefs faster, they would rely less on their prior beliefs on this level. Accordingly, the neural representation of the corresponding prediction regressor would be attenuated in high trait anxiety. The increased beta amplitude response to the prediction regressor in HTA corresponds to the finding for state anxiety in our previous work (Hein and Herrojo Ruiz, 2022), yet that outcome was aligned with poor belief updating and rigid prior beliefs in those individuals. Earlier work has found an association between alpha and beta activity and the encoding of predictions, which could contribute to the dampening of precision weights during perceptual processing (Auksztulewicz et al., 2017; Bauer et al., 2014; Sedley et al., 2016) and the suppression of model updates during visuomotor learning (Tan et al., 2016). These previous findings make the interpretation of the prediction results in the current study challenging. In addition, the interpretation is further limited because the prediction analysis was exploratory and the statistical results uncorrected. Further work is therefore needed to gain insight into the neural oscillatory representation of predictions in reward-based learning in anxiety. Invasive local field potential recordings across cortical and subcortical regions, when available, could help clarify this aspect.

An additional consideration is that the DKT anatomical atlas we used to obtain the surface-based cortical parcellations has 68 regions of interest, yet a finer spatial resolution would be relevant to identify and dissociate specific effects in some cortical regions, such as the vmPFC, or in subcortical regions such as the amygdala. An exciting prospect for follow-up studies is to use subject-specific 3D printed head-casts in MEG research to improve the signal-to-noise ratio and reduce the localisation error in MEG research (Bonaiuto et al, 2021).

## Conclusion

In summary, our findings provide important insights into the oscillatory neural correlates of the effect of higher-order environmental statistics on reward-learning behaviour in anxiety. Overall, the neural findings demonstrated that oscillatory signals in the ACC reflect a consistent modulation of belief updating and precision weighting by trait anxiety. The effects of anxiety on belief updating are also expressed in gamma changes in the dmPFC and lOFC, with additional involvement of the lOFC in encoding discrete lose outcomes. By identifying the source-level oscillatory correlates of Bayesian PC in subclinical trait anxiety, our study opens new prospects for neuromodulatory and neurofeedback interventions and pharmacological treatment in anxiety disorders.

## Materials and Methods

### Participants

We recruited 39 participants (24 female, 15 male) aged between 18 and 36 years (mean 22.8, SEM 0.9) who completed the MEG and behavioural study. We additionally acquired individual T1-weighted anatomical magnetic resonance images (MRI, details below). All participants reported having normal or corrected-to-normal vision. Individuals were excluded if they had a history of psychiatric or neurological disease or head injury, and/or were on medication for anxiety or depression. Written informed consent was obtained from all participants before the experiment, and the experimental protocol was approved by the ethics committee of the Institutional Review Board of the National Research University Higher School of Economics in Moscow, Russia.

Our sample size was estimated using the behavioural and EEG data from our recent work on decision making in state anxiety (Hein et al., 2020, 2021). MATLAB function sampsizepwr (two-tailed t-test) was used to estimate the minimum sample size for a statistical power of 0.80, with an a of 0.05. This function was evaluated on the HGF model parameter ω_2_ (the low-level tonic log-volatility estimate) and the beta activity modulation to pwPE, resulting in a minimum of 16 participants in each group (high, low anxiety). In the current MEG study, we recruited 20 and 19 participants in the LTA and HTA groups, respectively.

### Assessment of anxiety

Participants’ trait anxiety level was measured twice using Spielberger’s State-Trait Anxiety Inventory (STAI, trait subscale X2, 20 itemts, score 0–80; Spielberger 1983): one assessment prior to attending the experiment as a selection procedure, and one at the beginning of the experimental session (to validate the pre-screened level). Trait anxiety refers to a relatively stable metric of an individual’s anxiety level derived from the self-reported frequency of anxiety from past experiences (Grupe and Nitschke 2013). Trait anxiety in subclinical populations is commonly measured using the STAI trait subscale, a measure thought to reflect the general risk factor for an anxiety or affective disorder (Grupe and Nitschke 2013). This scale taps into the overall exaggerated perspective of the world as threatening, providing a good measure of how frequently a person has experienced anxiety across their life (Raymond et al. 2017).

We used the trait anxiety scores as a selection process to form the two experimental groups: low trait anxiety (LTA, defined as a STAI score below or equal to 36) and high trait anxiety (HTA, defined as a STAI score above 45). These values were selected to include the normative mean value in the working adult population as upper threshold in the LTA group (36, SD 9, Spielberger et al. 1983). In addition, the HTA threshold value was informed by the cut-off point (> 45) used to denote clinically significant anxiety in treatment studies in anxiety disorder patients (Fischer and Durham, 1999; See also Shadli et al., 2021). Trait anxiety scores ranged between 24 and 65. The average anxiety scores for each group were 30.5 (LTA, SEM 0.8) and 51.7 (HTA, SEM 1.5), comparable to LTA/HTA group values in recent investigations of reversal learning in trait anxiety (Xia et al., 2021; Jiang et al., 2018). Importantly, the experimental groups were balanced in terms of age and sex. The high trait anxiety group (HTA, mean age 22.6, SEM = 1.1) consisted of 12 females, while the low trait anxiety group (LTA, mean age 23.7, SEM = 1.0) consisted of 12 females. In addition to the trait inventory, measures of self-reported state anxiety using the STAI state subscale (X1, 20 items, score 0–80) were taken prior to the experiment and after completing the experiment.

In our previous work, we assessed heart rate variability (HRV) as a proxy measure for state anxiety (Hein et al. 2021). We calculated the coefficient of variation (CV = standard deviation/mean) of the difference intervals between consecutive R-peaks (inter-beat interval, IBI) extracted from the continuous electrocardiography (ECG) data as a metric of HRV. In previous empirical studies, states of anxiety have been shown to lower HRV and high-frequency HRV (HF-HRV, 0.15–0.4 Hz, see (Thayer et al. 1996; Hein et al. 2021; Sporn et al. 2020; Gorman and Sloan 2000; Friedman 2007). However, experiments on the association between trait anxiety levels and HRV/HF-HRV metrics have reported both reductions (Miu et al. 2009; Bleil et al. 2008; Mujica-Parodi et al. 2009) and small or inverse effects (Dishman et al. 2000; Narita et al. 2007). Thus, the relationship between trait anxiety and HRV remains unclear. In this study, we include a complimentary analysis of both HRV and HF-HRV to supplement our self-report measures of trait and state anxiety. See section on **Acquisition and preprocessing of MEG and ECG data** for further details.

### Experimental design and task

We used a between-subject experimental design with two anxiety groups: HTA and LTA. Participants performed a probabilistic binary reward-based learning task in a volatile learning setting (Behrens et al. 2007; de Berker et al. 2016; Iglesias et al. 2013; **Figure 1**). The session was split between an initial resting state block (R1: baseline) of five minutes and two experimental reward-learning task blocks consisting of a total 320 trials (TB1, 160 trials – TB2, 160 trials). During the continuous recording of MEG and ECG responses in the baseline block, participants were told to try to relax and fixate on a central point of the screen with their eyes open.

Similarly to Hein et al. (2021), participants were informed that the total sum of all their rewarded points would translate into a monetary reward at the end of the experiment. The calculation for this remuneration was the sum total of winning points divided by six plus 400, given in Russian rubles ₽ (for example, 960 points pays 960/6+400 = 560₽).

For every trial, a blue and an orange stimulus were shown on the monitor. Their location was either to the right or left of the centre, randomly generated in each trial. The maximum time allowed for a response before the trial timed out was 1300 ms ± 125 ms. In contrast to our previous EEG study (Hein et al. 2021; Hein and Ruiz 2021), responses here were given by pressing a button in a response box with either the left or right thumb (corresponding to selecting either the left or right image). After the participant made their choice, the selected image was outlined in bright green for 1000 ms (± 200 ms) to indicate their response. After, feedback of the trial outcome was provided (win, green; lose or no response, red) in the centre of the screen for 1250 ms (± 250 ms). To conclude a trial, a fixation cross was shown in the centre of the screen (1750 ms [± 250 ms]). Participants were told to select the image they believed would reward them to maximise reward across the 320 trials, and also to modify their selections in response to any inferred changes to their underlying probability. Prior to starting the experimental task blocks (TB1, TB2), each participant performed 16 practice trials and filled out the first state anxiety report. Between the two experimental task blocks, participants rested for a short self-timed interval. After completing the second task block, participants filled out the second state anxiety report before finishing the experiment.

### Modelling behaviour: The Hierarchical Gaussian Filter

To model behaviour, we used the Hierarchical Gaussian Filter (HGF, Mathys et al. 2011; Mathys et al. 2014, version 6.0.0, open-source software available in TAPAS, http://www.translationalneuromodeling.org/tapas, see Frässle et al. 2021). This model has been used widely to describe task responses in multiple learning contexts (de Berker et al. 2016; Iglesias et al. 2013; Marshall et al. 2016; Reed et al. 2020; Diaconescu et al. 2014; Weber et al. 2020; Suthaharan et al. 2021). We used TAPAS in Matlab R2020b.

As in our previous work (Hein et al. 2021), we utilised a generative perceptual model for binary outcomes termed the 3-level HGF (Mathys et al. 2011). The input to the model was the series of 320 outcomes and the participant’s responses. Observed outcomes in trial *k* were either u^(k)^ = 1 if the blue image was rewarded (orange stimulus unrewarded) or u^(k)^ = 0 if the blue stimulus was unrewarded (orange stimulus rewarded). Trial responses were defined as y^(k)^ = 1 if participants chose the blue image, while y^(k)^ = 0 corresponded to the choice of the orange image. In the 3-level HGF, the first level x_1_^(k)^ represents the true binary outcome in a trial *k* (either blue or orange wins) and beliefs on this level feature expected (irreducible) uncertainty due to the probabilistic nature of the rewarded outcome (Soltani and Izquierdo, 2019). In the absence of observation noise, u^(k)^ = x_1_^(k)^. The second level x_2_^(k)^ represents the true tendency for either image (blue, orange) to be rewarding. And the third level x_3_^(k)^ represents the log-volatility or rate of change of reward tendencies (Bland and Schaefer, 2012; Yu and Dayan, 2005). In the HGF update equations, the second and third level states, x_2_^(k)^ and x_3_^(k)^, are modelled as continuous variables evolving as Gaussian random walks coupled through their variance (inverse precision).

The coupling function between levels 2 and 3 is as follows (dropping index *k* for simplicity):

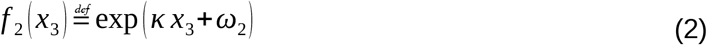

In equation (2), ω_2_ represents the invariant (tonic) portion of the log-volatility of x_2_ and captures the size of each individual’s stimulus-outcome belief update independent of x_3_. The κ (*Kappa*) parameter establishes the strength of the coupling between x_2_ and x_3_, and thus the degree to which estimated environmental volatility impacts the learning rate about the stimulus-outcome probabilities—here κ was fixed to one as in previous work (de Berker et al. 2016; Weber et al. 2020; Hein et al. 2021). On the other hand, the step size of x_3_ depends on the exponential of a positive constant parameter ω_3_ (the lower ω_3_ the slower participants update their beliefs about volatility). As in our prior work (Hein et al., 2021), parameters ω_2_ and ω_3_ were estimated in each individual (3-level HGF and HGF*_μ_*_3_; for the 2-level HGF, ω_3_ was fixed; **Table S1**).

We paired the 3-level HGF perceptual model with two alternative response models that map a participant’s beliefs to their decisions using a unit-square sigmoid function: i) with a fixed parameter *ζ* that can be interpreted as inverse decision noise that shapes choice probability: the sigmoid function approaches a step function as *ζ* tends to infinity (for further detail see Eq. 18 in Mathys et al. [2011]) ii) where the inverse decision noise is a function of the prediction of log-volatility: 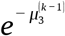, thus depending on the participant’s trial-wise beliefs on volatility—termed hereafter HGF*_μ_*_3_ (Diaconescu et al. 2014). The response model parameter ζ was also estimated in the 3-level and 2-level HGF models, while parameters *μ_3_^(0)^* and σ_3_^(0)^ were estimated in model HGF*_μ_*_3_ (**Table S1**). Simulations conducted to assess the accuracy of parameter estimation in the HGF models demonstrated that the most accurate estimation was for parameters ω_2_ and *μ_3_^(0)^*, while ω_3_ was poorly recovered, as shown previously (Reed et al., 2020; Hein et al., 2021; **Supplementary file 3**).

We direct the reader to the original methods papers for more details on the derivation of the perceptual model and equations of the HGF quantities used in this paper (Mathys et al. 2011; Diaconescu et al. 2014). Using the prior parameter values (**Table S1**) and series of inputs, maximum-a-posteriori (MAP) estimates of model parameters were then quantified and optimised using the quasi-Newton optimisation algorithm (Diaconescu et al. 2014; Cole et al. 2020; Reed et al. 2020).

Participants’ decisions may best be explained by simpler non-hierarchical non-Bayesian models. To address this, we compared a Bayesian family of models to a family of reinforcement learning models, as in previous work (Lawson et al. 2020; Cole et al. 2020; de Berker et al. 2016). For the Bayesian family, we included three models. The first was the binary 3-level HGF (HGF_3_) that captures volatility estimates and uses a set decision noise parameter for mapping beliefs to decisions (Mathys et al. 2011). The second was the 3-level HGF where decisions depend dynamically on estimated volatility (HGF*_μ_*_3_, Diaconescu et al. 2014). In addition, we used a version of the HGF with only two levels (HGF_2_) where the third level volatility is fixed. For the family of reinforcement learning models, we used a reward-maximising Rescorla-Wagner (RW) model with a fixed learning rate (see (Rescorla and Wagner 1972) for further details) and a Sutton K1 model (SK1) using a dynamic learning rate (Sutton 1992). Model comparison was performed using random-effects Bayesian model selection (BMS, see Stephan et al. 2009) using code from the MACS toolbox (Soch and Allefeld 2018).

### Acquisition and preprocessing of MEG and ECG data

Neuromagnetic brain activity was recorded using a 306-channel MEG system (102 magnetometers and 204 gradiometers, Elekta Neuromag VectorView, Helsinki, Finland) in sitting position. We used a head-position indicator to control for head movements, with four coils affixed to the head, two placed on the top of each side of the forehead, and two on the mastoid process of each side. Eye movements were controlled using an electrooculogram (EOG): Two horizontal EOG electrodes were placed each side of the temple, while the two vertical EOG electrodes were placed above and below one eye. In addition, two electrodes were used for electrocardiography (ECG) recording using in a two-lead configuration montage (Moody and Mark 1982). MEG, EOG, and ECG signals were recorded with a sampling rate of 1000 Hz and a band-pass filter of 0.1–330 Hz. Following the MEG acquisition phase, we de-noised the signals and corrected head movements using the Temporally extended Signal-Space Separation (tSSS) method (Taulu and Hari, 2009), built-in in the Elekta software (Maxfilter^TM^; Elektra Neuroscience 2010; settings: sliding window = 10 s, subspace correlation threshold = 0.9).

Further preprocessing of the MEG data (magnetometers and planar gradiometers) was conducted with the MNE-python toolbox (Gramfort et al., 2013; Python version 3.9.4), as well as additional custom Python scripts. For the HRV/HF-HRV analyses, the ECG signal was pre-processed using the FieldTrip toolbox (Oostenveld et al. 2011) for MATLAB® (v. 2020b, The MathWorks, Natick, MA).

The MEG signals were downsampled to 250 Hz. Next, we removed power-line noise by applying a zero-phase notch filter at 50 Hz and 100 Hz and removed biological artefacts (eye movements, blinks, heartbeats) using independent components analysis (ICA, fastICA algorithm). MEG signals that exceeded a certain amplitude threshold (5^-12^ T for magnetometers, 4^-10^ T/cm for gradiometers) were excluded from further analysis. We also used the standard MNE-python algorithm for automatic detection of ICs relating to EOG and ECG artifacts, which were however validated visually in each subject. On average, we removed 4.5 components (SEM 0.1).

During preprocessing of the ECG signal, we extracted cardiac events (the QRS-complex, R wave peak) using the FieldTrip toolbox. Afterwards, we calculated the latency of each R-peak and the CV of the inter-beat interval (IBI), our proxy metric for HRV across both experimental task blocks. The CV for each participant was normalised to the resting state block (R1: baseline). To estimate the high-frequency content of the HRV (HF-HRV) we used the inter-beat-interval (IBI) time series. First, we interpolated at 1Hz with a spline function (order 3), and subsequently estimated spectral power using Welch’s periodogram method (Hanning window, following Rebollo et al. 2018). Estimates of power were normalised to the average power in R1 and converted to decibels (dB) for statistical analysis.

### Structural Magneto Resonance Imaging

Structural brain MRIs (1 mm3 T1-weighted) were obtained for all participants and used for source reconstruction. The MRI image was derived from a 1.5T Optima MR 360 system (Spin Echo sequence, slice thickness 1 mm, field of view 288 x 288, TR = 600, TE = 13.5).

### Source Analysis

Source localisation of the MEG signals was performed using Linearly Constrained Minimum Variance beamformers (Van Veen et al., 1997) in MNE-Python (Gramfort et al., 2013). First, we used the individual T1-weighted MRI images to construct automatic surface-based cortical parcellations in each hemisphere with Freesurfer 6.0 software (http://surfer.nmr.mgh.harvard.edu/; Dale, Fischl, & Sereno, 1999; Fischl, Sereno, & Dale, 1999). We chose the label map of the Desikan–Killiany–Tourville atlas (DKT, Desikan et al. 2006), which parcellates the cerebral cortex into 68 regions of interest (ROIs). Subcortical parcellations were also generated as default in Freesurfer but were not used in this study. Coregistration of the MR and MEG coordinate systems was performed with an automated algorithm in MNE-python available in the MNE software (mne_analyze: http://www.martinos.org/mne/stable/index.html). The coregistration step used the HPIs and the digitized points on the head surface (Fastrak Polhemus). We additionally verified that the coregistration of three anatomical (fiducial) locations (the left and right preauricular points and the nasion) were correct in both coordinate systems.

For forward model calculations, we used the command-line tool “mne watershed” to compute boundary element conductivity models (BEM) for each participant and selected the inner skull surface as volume conductor geometry. Then, we created a surface-based source space with “oct6” resolution, leading to 4098 locations (vertices) per hemisphere with an average nearest-neighbor distance of 4.9 mm.

For inverse calculations, LCMV beamformers were used. The adaptive spatial filters were computed with a data-covariance matrix in the target interval (0–1.8 s in outcome-locked and stimulus-locked analyses) and a noise-covariance matrix in a time interval preceding the stimulus (−1 to 0 s pre-stimulus) and outcome events (−3 to −2 s pre-outcome, thus corresponding to a waiting period before the stimulus). The regularization parameter λ was set to 5%. Prior to LCMV beamforming, the MEG signals were band-pass filtered between 1 and 40 Hz for the alpha and beta range and between 30 and 124 Hz for the gamma range (below the Nyquist rate at 125 Hz). Last, source estimate time courses for individual vertices were obtained for a set of cortical labels corresponding to our ROIs: anterior cingulate cortex (rostral ACC [rACC], caudal ACC [cACC]), medial OFC and lateral OFC, which include the vmPFC according to previous MEG work (Yuan et al., 2020, but see Rushworth et al., 2011 for a debate on the vmPFC delineation), dorsomedial PFC (dmPFC; label superior frontal gyrus, SFG). The representative time course per label was obtained using the “PCA flip” method in MNE-Python. This method consists of applying singular value decomposition to each vertex-related time course in the label, followed by extraction of the first right singular vector. Next, each vertex’s time course is scaled and sign flipped. Following this procedure, we obtained five bilateral (10 in total) time courses corresponding with our three ROIs. An additional exploratory analysis was carried out in the other labels of the DKT atlas to identify effects outside of our ROIs.

### Spectral Analysis and Convolution Modelling

We estimated standard time-frequency representations of the source-level time series using Morlet wavelets. TF spectral power was extracted between 8 and 90 Hz. For alpha (8–12 Hz) and beta (13–30 Hz) frequency ranges we used 5–cycle wavelets shifted every sampled point in bins of 2 Hz (Kilner et al. 2005; Litvak et al. 2011; Ruiz et al. 2009). For gamma-band activity (32–100 Hz), 7-cycle wavelets sampled in steps of 2 Hz were used.

After transforming the source-level time series to TF representations, we used linear convolution modelling for oscillatory responses (Litvak et al. 2013). This analysis was implemented in SPM 12 (http://www.fil.ion.ucl.ac.uk/spm/) by adapting code developed by Spitzer et al. (2016) freely available at https://github.com/bernspitz/convolution-models-MEEG. This method allowed us to model the pseudo-continuous TF data resulting from concatenated epochs as a linear combination of explanatory variables (parametric HGF regressors or discrete stimulus, response and outcome regressors) and residual noise. The general linear model explains this linear combination as follows:

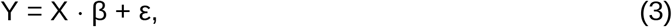

here *Y* ∈ ℝ^t*x*f^ denotes the measured signal, the TF transformation of the pseudo-continuous time series, and is defined over *t* time bins (trials x peri-event bins in our study) and *f* frequencies. The linear combination of *n* explanatory variables or regressors is defined in matrix *X*∈ℝ^txn^, and modulated by the regression coefficients *β*∈ℝ^nx*f*^. The noise matrix is denoted by ε∈ℝ^tx*f*^. Matrix *X* is specified as the convolution of an input function, encoding the presence and value of discrete or parametric events for each regressor and time bin, and a Fourier basis function. This problem is solved by finding TF images R_i_ for a specific type of event *i* (e.g. outcome or response event type):

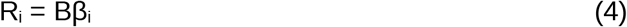

In the expression above, B denotes a family of *m* basis functions (sines, cosines) used to create the regressor variables *X* by convolving the basis functions B∈ℝ^pxm^ with *k* input functions U representing the events of interest at their onset latencies (U∈ℝ^txk^), and thus X = UB. Using ordinary or weighted least squares, the predictors *β_i_* are estimated over frequences and basis functions for each regressor *i*. The TF response images R_i_ ∈ℝ^px*f*^ have dimensions *p* (peri-event interval of interest) and *f*, and can be interpreted as deconvolved TF responses to the event types and associated parametric regressors. The TF images R_i_ can be used for standard statistical analysis. A schematic of the convolution modelling approach is presented in **Figure 6**.

**Figure 6.**
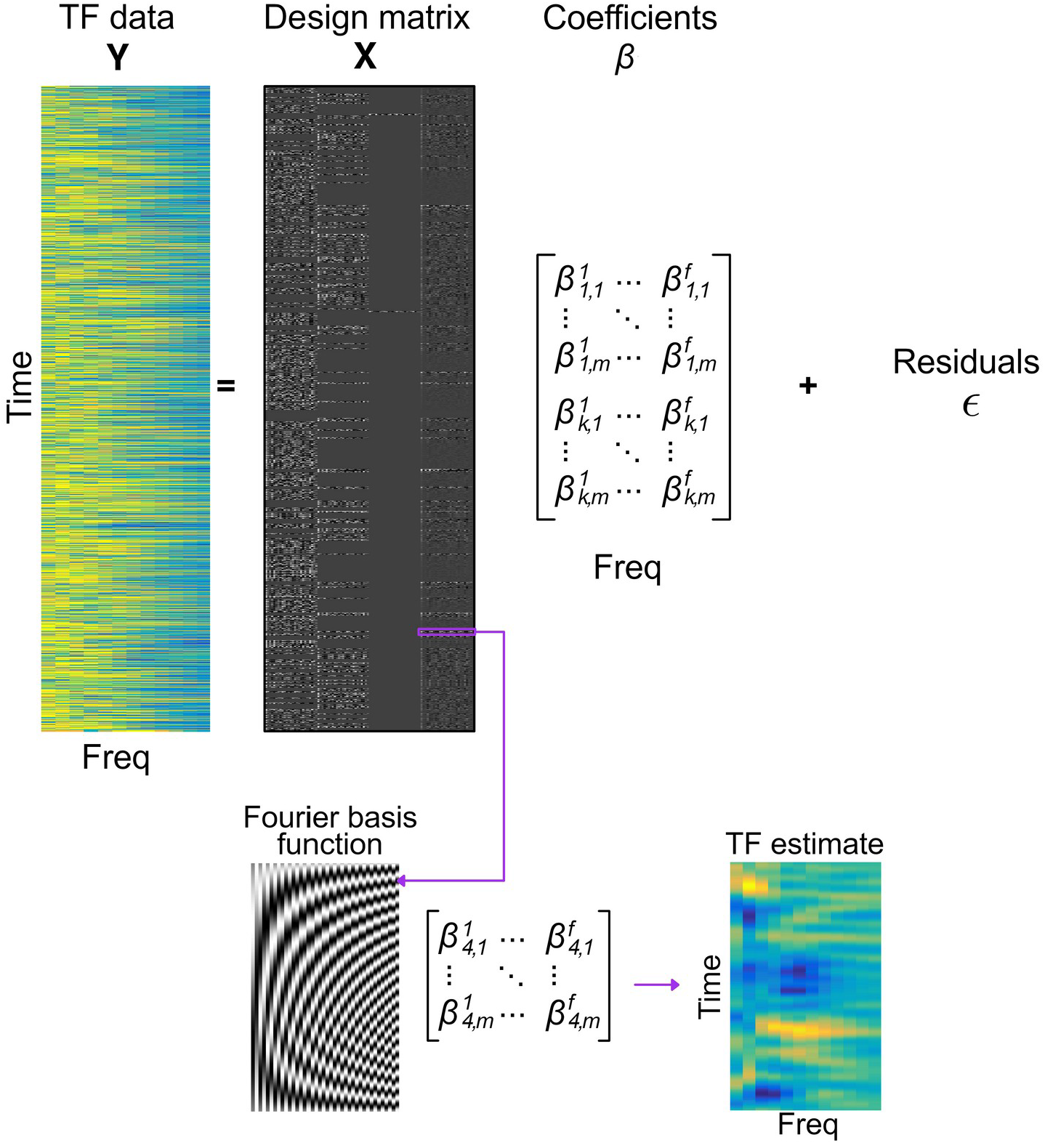
Convolution general linear model. Standard pseudo-continuous time-frequency (TF) representations of the source-level MEG signal (Y) were estimated using Morlet wavelets. In GLM, signals Y are explained by a linear combination of explanatory variables or regressors in matrix X, modulated by the regression coefficients β, and with an added noise term (ɛ). The design matrix X shown in this figure was constructed by convolving Fourier functions (*m* = 20; 20 sine, 20 cosine functions; left inset at the bottom) with a set of input functions representing the onsets and value of relevant discrete or parametric events. In this example, we used *k* = 4 regressors (columns left to right): Outcome Win, Outcome Lose, Outcome No Response, and absolute pwPE on level 2, which were defined over time. Solving a convolution GLM provides *response images* (TF estimate in the figure, arbitrary unit) that are the combination of the basis functions *m* and the regression coefficients β_i_ for a particular regressor type *i* defined over frequencies *f* and basis functions *m*. Thus, convolution GLM effectively estimates deconvolved TF responses (TF estimate, rightmost image at the bottom) to the event types and associated parametric regressors.

To adhere to the GLM error assumptions (Kiebel et al. 2005; Litvak et al. 2013) we first converted the spectral power to amplitude by executing a square-root transformation. Our trial-wise explanatory variables included discrete regressors coding for stimuli (blue image left, blue image right), responses (right, left, no response), outcome (win, lose, no response) and relevant parametric HGF regressors: precision-weighted prediction errors (pwPEs) encoding the steps of the belief updating process (|ε_2_|, **Figure 2B**) and unsigned predictions about the tendency towards a stimulus-reward contingency on level 2 (| *μ̂*_2_|, hereinafter termed ‘predictions’). For computational efficiency, we conducted separate GLMs for outcome-locked and stimulus-locked analyses, inserting the relevant discrete and parametric regressors at the corresponding latencies in each case (see **Results**).

In our main GLM assessing pwPEs, similarly to Hein et al. (2021), we found high linear correlations between the absolute value of the second-level pwPEs, |ε_2_|, and the third-level pwPEs about environmental change (ε_3_; the Pearson correlation coefficients ranged from 0.67 to 0.95 among all 39 participants). Due to multicollinearity of regressors, pwPEs on level 3 have been excluded from subsequent analysis (see Vanhove 2021; Hein et al. 2021). We conducted a separate convolution model to assess the dissociable effect of the precision weights (σ_2_) and unsigned PE (|δ_1_|) regressors—which combined represent the unsigned pwPEs on level 2—on the TF responses. In our exploratory analysis of the neural correlates of predictions, we choose the absolute values of predictions on level 2 | *μ̂*_2_|, and excluded the third level log-volatility predictions *μ̂*_3_. Similarly to the pwPE GLM, this decision was grounded on multicollinearity of regressors: There were high linear correlations between |*μ̂*_2_| and log-volatility *μ̂*_3_ (Pearson r between −0.95 and 0.37, N = 39).

Our primary convolution model aimed to assess the parametric effect of pwPEs about stimulus outcomes on TF responses in 8-90Hz in a relevant time interval following the outcome event. The impulse response design matrix thus included as parametric regressor the pwPE values at the latency of the outcome regressor and the outcome events as discrete regressors. This model was estimated using a window from −500 to 1800 ms relative to the outcome event (“outcome-locked” analysis), informed by our previous work in state anxiety (Hein and Ruiz 2021). This window was refined during the subsequent statistical analysis (see below). Our second GLM evaluated the separate effect of precision weights and surprise about stimulus outcomes on the alpha/beta oscillatory activity also from −500 to 1800 ms post-outcome. The input matrix included the outcome (win/lose/no response) events as discrete regressors, and parametric regressors σ_2_ and |δ_1_|. Last, in an exploratory analysis we estimated the effect of the parametric prediction regressor on the alpha/beta oscillatory activity from −500 to 1800 ms around the stimulus event (“stimulus-locked” analysis). This GLM included as additional discrete regressors, the stimuli (blue right, blue left) and response (press left, press right, no response) events. In all convolution analyses, each discrete and parametric regressor was convolved with a 20th-order Fourier basis set (40 basis functions, 20 sines and 20 cosines). This setting allowed the GLM to resolve modulations of TF responses up to ∼8.7 Hz (20 cycles / 2.3 seconds; or ∼115 ms). In an additional control analysis, we used a 40th-order Fourier basis set to assess gamma activity modulations by the unsigned pwPE regressor (**Figure 3 – figure supplement 3**). This set provided a temporal resolution of 57.5 ms.

### Statistical Analysis

Statistical analysis of standard behavioural and computational model variables focused on between-group contrasts (LTA, HTA). Our dependent variables (DVs) were i) win rates, win-stay/lose-shift rates, total switch rates, RT (ms); ii) CV and spectral measures (expressed in dB) of HRV, which were normalised to the baseline (R1) average values at rest; iii) HGF belief trajectories averaged across trials in each task block separately: a) informational uncertainty about the stimulus outcomes (σ_2_); b) mean of the posterior distribution of beliefs about volatility (μ_3_), and the associated posterior uncertainty (variance, σ_3_); c) environmental uncertainty: exp(κμ_3_^(k-1)^ + ω_2_), which is greater if the environment is more volatile; iv) HGF perceptual model parameter quantities ω_2_ and ω_3_. We conducted 2 x 2 Group x Block (TB1, TB2) factorial analyses in the following DVs: win rate, HRV/HF-HRV. This was implemented using non-parametric factorial synchronised rearrangements (Basso et al. 2007) with 5000 permutations. Between-group comparisons of DV iii-iv and i (win-stay/lose-shift rates, RT), were carried out using pair-wise permutation tests (5000 permutations).

To address the multiple comparisons problem, where it arises (e.g. several post-hoc analyses), we control the false discovery rate (FDR) using an adaptive linear step-up procedure set to a level of q = 0.05 providing an adapted threshold p-value (*P_FDR_*, Benjamini et al. 2006). In the case of pair-wise statistical analyses we provide estimates of the non-parametric effect sizes for pair-wise comparisons and associated bootstrapped confidence intervals (Grissom and Kim 2012; Ruscio and Mullen 2012). The within-group effect sizes are estimated as the probability of superiority for dependent samples (Δ*_dep_*), while the between-group effect sizes are based on the probability of superiority (Δ, see Grissom and Kim (2012).

Statistical analysis of the source-level TF responses obtained in convolution modelling was performed with the FieldTrip Toolbox (Oostenveld et al. 2011), after converting the SPM TF images (in arbitrary units, a.u.) to a Fieldtrip structure. Given the large inter-individual differences typically observed in the amplitudes of MEG neuromagnetic responses, the source-level TF images were baseline corrected by subtracting the average baseline level (−300 to −50 ms) and dividing by the baseline standard deviation (SD) of the interval. We used a cluster-based permutation approach (two-sided t-test, 1000 iterations; Maris and Oostenveld 2007; Oostenveld et al. 2011) to assess between-group differences in TF responses across 10 anatomical labels, time points, and frequency bins (8–100 Hz for the outcome-locked pwPE model; 8–30 Hz for the outcome-locked precision ratio and unsigned PE model; 8–30 Hz for the exploratory stimulus-locked precision model). We did not consider spatial relations between anatomical labels but focused on spectrotemporal clusters. Based on the latency of the effects in our previous work (Hein and Ruiz, 2022), we chose as the temporal intervals of interest for the statistical analysis 200–1800 ms for the outcome-locked convolution models, and 100–700 ms for the stimulus-locked GLM. This analysis controlled the family-wise error rate (FWER) at level 0.025.

## Acknowledgements

TPH was funded by the Economic and Social Research Council (ESRC) and the South East Network for Social Sciences (SeNSS) through grant ES/P00072X/1. This work was partially supported by the Basic Research Program of the National Research University Higher School of Economics (Russian Federation) and has been carried out using HSE unique equipment in 2021 (Reg. Num 354937).

## Competing interests

The authors declare having no financial or non-financial competing interests.

## Code and Data availability

Code for the source reconstruction analysis (MNE Python) and convolution modelling (Matlab / SPM) has been deposited in the Open Science Framework Data Repository under the accession code wsjgk. Behavioural and computational modelling data are also provided.

## Notes

**Conflict of interest statement:** The authors declare no competing financial interests

### Competing Interest Statement

The authors have declared no competing interest.

### Summary of Updates

The figures have been embedded in the text, as opposed to being appended at the end as in v1.

