## Supplementary Material for "Changes in oscillations in anterior cingulate and medial prefrontal cortex are associated with altered signatures of Bayesian predictive coding in trait anxiety"

### Figure Supplements

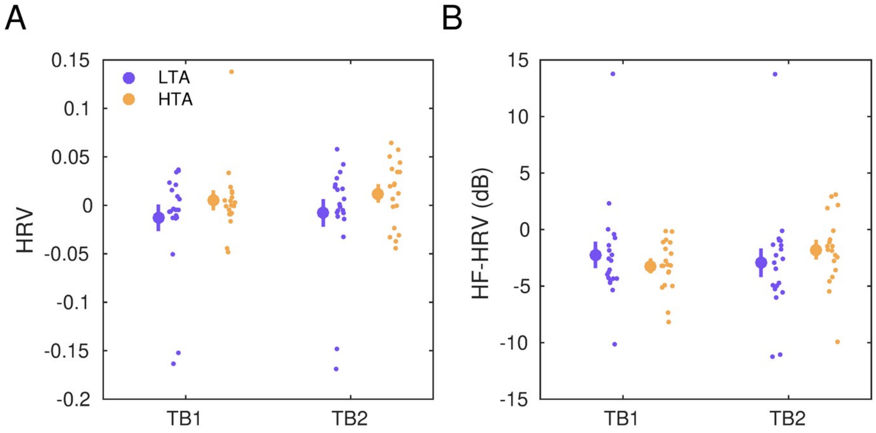


**Figure 1 – figure supplement 1. Heart-rate variability in trait anxiety. A)** Normalised heart-rate variability (HRV). Average HRV (coefficient of variation of the inter-beat-interval of the R-peak in the ECG signal) in high trait anxiety (HTA, yellow) and low trait anxiety (LTA, dark blue). The HRV from both experimental task blocks has been normalised by subtracting the average HRV in the resting state baseline (R1). No significant differences were found using a non-parametric 2 x 2 Block x Group factorial analysis with synchronised rearrangements ($P > 0.05$ for main and interaction effects). **B)** Normalised high-frequency (HF) HRV. Analysis of the high frequency (0.15 – 0.40 Hz) spectral content of the inter-beat-interval (IBI) time series data revealed there was no significant difference between HTA relative to LTA ($P > 0.05$ as in A).

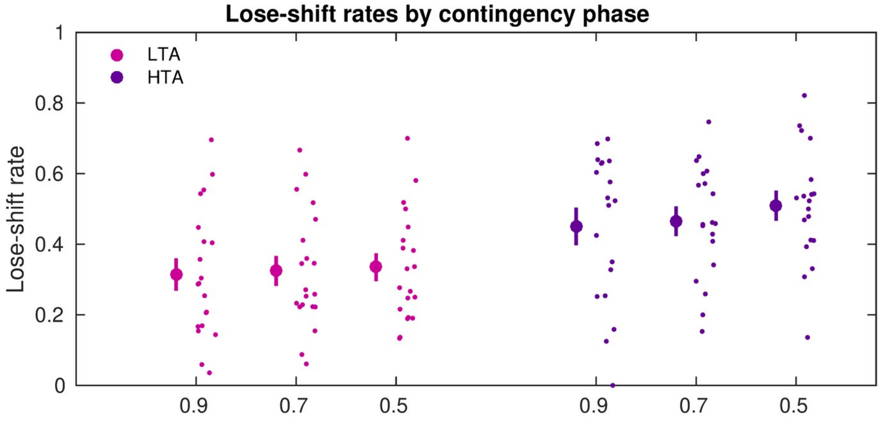


**Figure 1 – figure supplement 2.** Lose-shift rate across contingency phases. Illustration of the lose-shift rate, as in Figure 1D, across contingency phases: 0.9/0.1 and 0.1/0.9, 0.7/0.3 and 0.3/0.7, 0.5/0.5. Both groups (LTA: magenta; HTA: purple) of participants exhibited similar lose-shift rates across changes in stimulus-reward mappings.

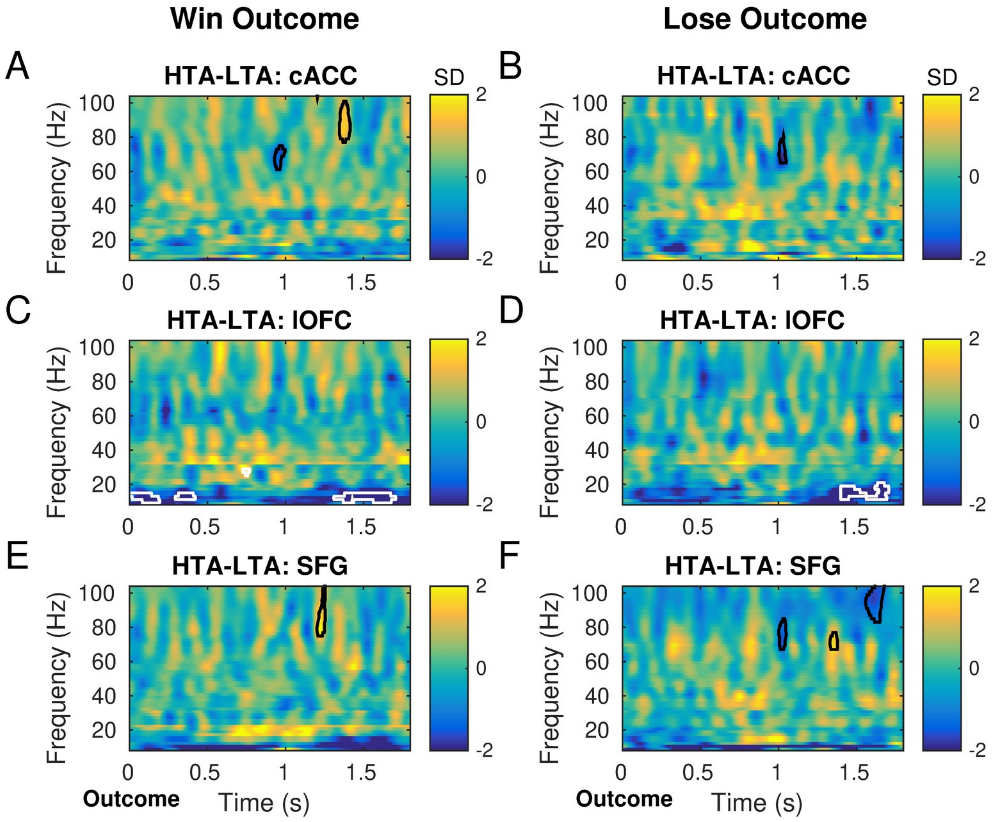


**Figure 3 – figure supplement 1. Effect of win and lose regressors on gamma and alpha/beta power amplitude in pwPE convolution model.** Panels **A, C, E** display the statistical results of the pwPE convolution model for the Win Outcome regressor. Between-group differences are denoted by the white and black contours (Significant clusters are FWER-controlled); Panels **B, D, F,** same as A, C, E but for the Lose Outcome regressor. See main text.

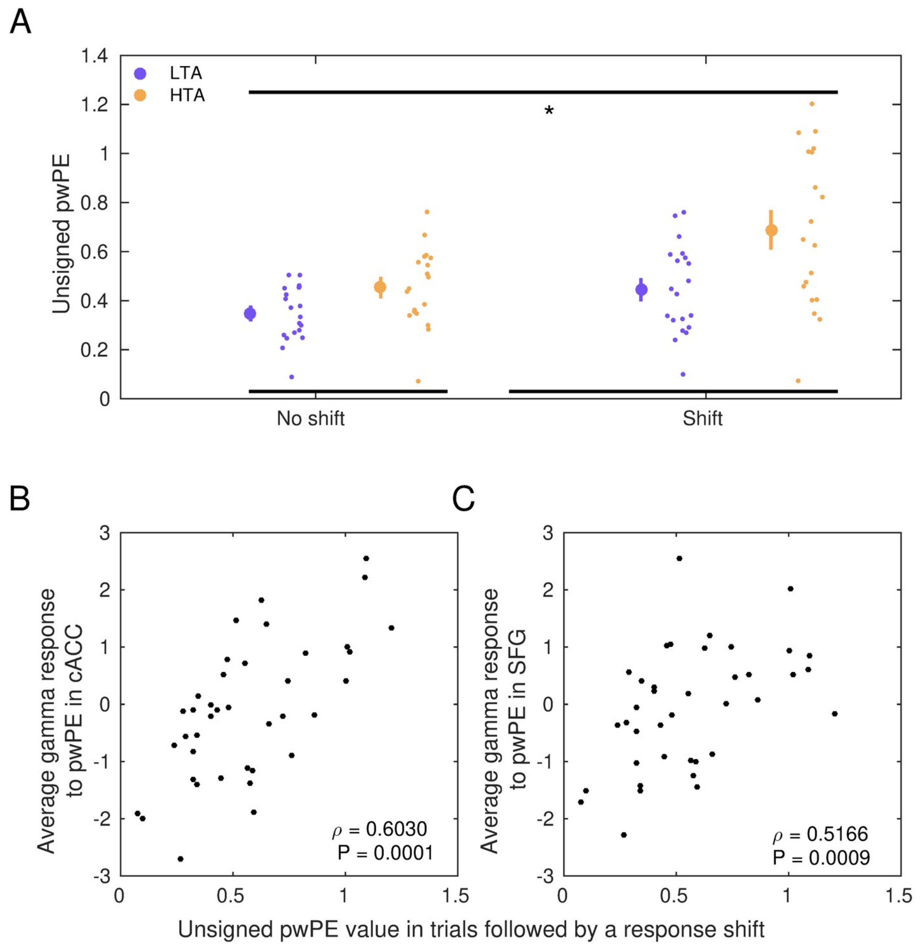


**Figure 3 - figure supplement 2. Trials leading to a shift in response choice are associated with larger unsigned pwPE values. A)** The values of unsigned pwPE updating of beliefs about stimulus-reward contingencies (denoted by $|\varepsilon_2|$) were larger in trials that were followed by a response shift (significant main effect of factor Shift, 2 x 2 synchronised rearrangements, $P$ = 0.0012; denoted by the black line on top and the asterisk). High relative to low trait anxiety individuals also had larger $|\varepsilon_2|$ overall (main effect of Group, $P$ = 0.0068; denoted by the black lines at the bottom) and a more pronounced dissociation between the $|\varepsilon_2|$ values in trials followed by a response shift or repetition (significant interaction, $P$ = 0.0200). Data in each group are represented using the average (large dot) with SEM bars. To the right are individual data points to display dispersion. HTA, high trait anxiety (yellow); LTA, low trait anxiety (purple). **B)** Non-parametric correlation between the average gamma response to the unsigned pwPE regressor in the cACC and the $|\varepsilon_2|$ values in trials followed by a response shift (Spearman ρ = 0.6030, P = 0.0001; the gamma average was estimated in the significant cluster of between-group differences, **Figure 3**). **C)** same as B) but for the dmPFC (ρ = 0.5166, P = 0.0009).

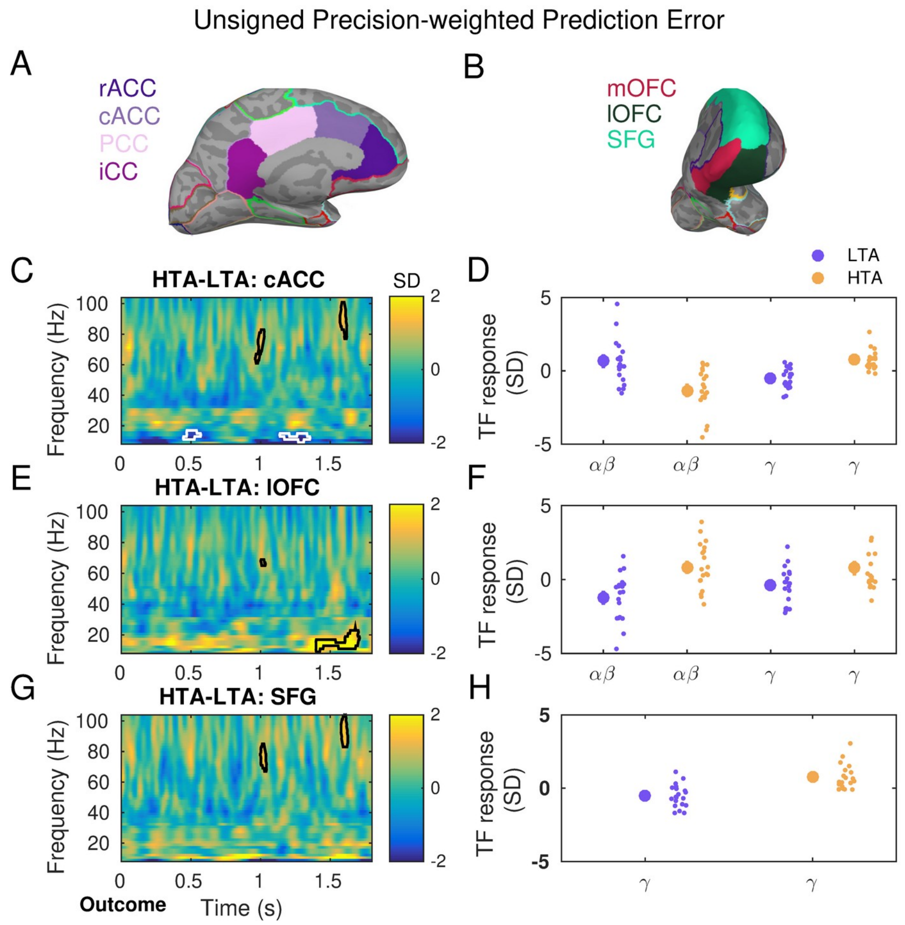


**Figure 3 - figure supplement 3.** Same as **Figure 3**, but using a 40th-order Fourier basis set for the gamma-band convolution model. Between-group differences are denoted by the white and black contours (Significant clusters are FWER-controlled). All cluster effects extend for at least one cycle of the associated frequency.

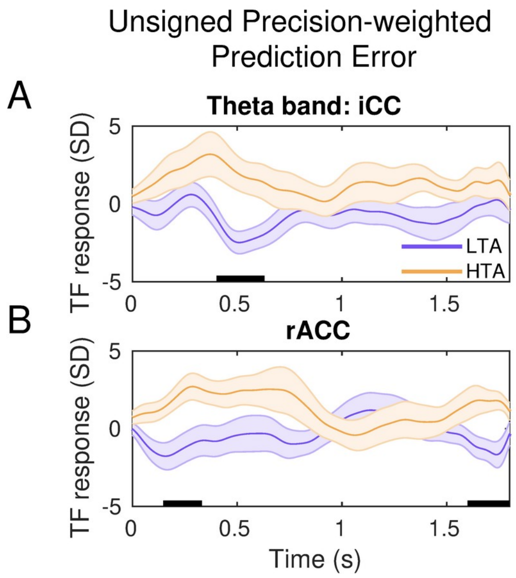


**Figure 3 - figure supplement 4.** Same as **Figure 3**, but running the convolution model in the theta (4–7 Hz) frequency range in an exploratory analysis. Between-group significant differences in TF responses to unsigned precision-weighted PEs (FWER-controlled). HTA, high trait anxiety (yellow); LTA, low trait anxiety (purple).

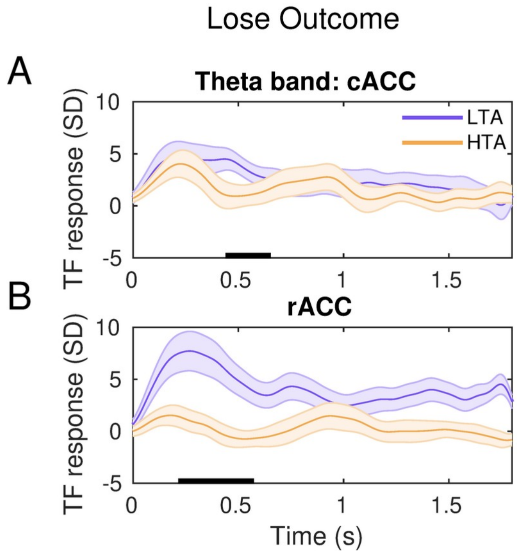


**Figure 3 - figure supplement 5.** Same as **Figure 3**, but running the convolution model in the theta (4–7 Hz) frequency range. Between-group significant differences in TF responses to the discrete Lose Outcome regressor (FWER-controlled). HTA, high trait anxiety (yellow); LTA, low trait anxiety (purple).

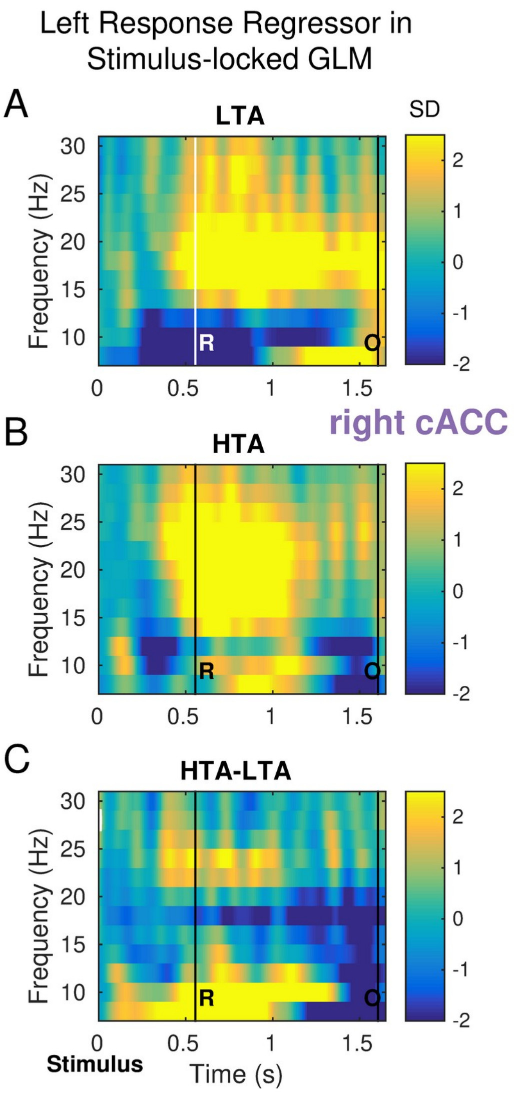


**Figure 5 Supplement figure 1. Stimulus-locked analysis of the response regressor. A-B)** Same as panels A and B in **Figure 5. C)** No significant between-group differences in the TF images to the discrete regressors were observed in any of our ROIs or additional anatomical labels ($P > 0.05$).

**Supplementary file 1**

**Measures of anxiety**

Factorial testing on the normalised HRV index was conducted usinga non-parametric 2 x 2 Group x Block factorial analysis with synchronised rearrangements. The results revealed a non-significant main effect of Group (*P* = 0.57), Block (*P* = 0.21) and interaction effect (*P* = 0.91, see **Figure 1 – figure supplement 1A**). Analysis of the spectral characteristics of the IBI time series using a non-parametric 2 x 2 factorial test also demonstrated no significant main effect of Group (*P* = 0.78), Block (*P* = 0.68) and interaction effect (*P* = 0.08) on HF-HRV (**Figure 1 – figure supplement 1B**).

Subjective self-reported measures of state anxiety showed a significant main effect of the Group factor (HTA: mean 35.4, SEM 1.9; LTA: mean 27.9, SEM 1.1, *P* < 0.0001). There was no effect of the factor Block (*P* = 0.23) or interaction effect (*P* = 0.22). This analysis demonstrates that the HTA participants were subjectively more anxious than LTA during both task blocks. However, the state anxiety scores in each group were relatively smaller than the trait scores, which suggests that task performance did not induce high levels of sustained anxiety.

**Supplementary file 2**

### Reaction time

Averaged reaction times (RT) were assessed in seconds (s). HTA participants made their decisions with an average RT of 0.493 (SEM 0.010) s, while LTA participants made a choice at 0.495 (SEM 0.017) s. We found no significant main effect of the Group factor and a non-significant interaction effect ($P$ = 0.2700, $P$ = 0.4400, respectively), consistent with our previous work in Hein et al. (2021) and prior studies in anxiety (Bishop 2009). In contrast to the findings in our previous study (Hein et al. 2021), however, there was no significant effect of the factor Block ($P$ = 0.23). Next, we asked whether RT in lose and win trials differed statistically, after collapsing the group information (joint sample of all participants). Unlike in Hein et al., (2021), we did not observe significant differences in the RT of win (0.491 [0.009] s or lose trials (0.500 [0.011] s; $P$ = 0.540).

Last, we assessed whether classical attentional effects account for between-group behavioural differences. Anxiety-related attentional deficits during the task would be expressed by increases in RT in the predictable (0.9/0.1 and 01/0.9) but not in the unpredictable (0.5/0.5) phases of the task. To test this, we first averaged the RT values independently in the phases with 0.5/0.5 and 0.9/0.1 – 0.1/0.9 probabilistic mappings. Next, we run planned pair-wise permutation tests. These assessed the mean RT between HTA and LTA groups independently for each probabilistic mapping. For the 0.9/0.1 – 0.1/0.9 contingency phase, there was no significant difference between RT in HTA (mean 496, SEM 16) and LTA (mean 513, SEM 18, $P$ = 0.41). Similarly, no difference was found for the 0.5–0.5 contingency phase between RT in HTA (mean 497, SEM 17) and LTA (mean 523, SEM 21, $P$ = 0.38). The results above are both consistent with Hein et al. (2021) and can be interpreted as evidence that no deficit in classical attentional resources was driving poorer task performance in our HTA group (Prinzmetal et al. 2009).

**Supplementary file 3**

We used simulations to evaluate the reliability of our estimates for the free model parameters in our implementation of the best fitting HGF model ($HGF_{\mu 3}$). In this model, the parameters that were estimated in each individual were $\omega_2$, $\omega_3$, $\mu_3^{(0)}$ and $\sigma_3^{(0)}$ (**Table S1**).

We simulated behavioural responses of 100 agents for six different values of $\omega_2$ (total 600 simulations), and seven different values of $\omega_3$ (total 700), when observing the input of one of our participants (ID zhanx_25). To determine the accuracy of the estimation of parameter $\mu_3^{(0)}$, we conducted similar simulations in 100 agents for six values of $\mu_3^{(0)}$.

These analyses were implemented using function tapas_simModel.m of the HGF toolbox (loop on om2 and om3, representing $\omega_2$, $\omega_3$, respectively; and loop on the number it iterations, N = 100):

```
sim = tapas_simModel(u, 'tapas_hgf_binary',  [NaN 0 1 NaN 1 1 NaN 0 0 1 1 NaN om2 om3],
'tapas_unitsq_sgm_mu3',123456789);
```

The simulated behavioural responses sim.y and the input u observed by participant #3 were then fitted with the tapas_fitModel.m function, similarly to the way we fitted the standard empirical data in our paritcipants:

```
est = tapas_fitModel(sim.y, u, hgf_binary_config, unitsq_sgm_mu3_config, optim_config)
```

with

```
optim_config = tapas_quasinewton_optim_config()
unitsq_sgm_mu3_config = tapas_unitsq_sgm_mu3_config()
hgf_binary_config = tapas_hgf_binary_config()
```

Note that in the above analyses the prior values of model parameters and initial values of the belief trajectories were modified in tapas_hgf_binary_config.m to correspond to the prior values for model $HGF_{\mu 3}$ given in **Table S1.**

This analysis demonstrated high accuracy for estimating $\omega_2$ and $\mu_3^{(0)}$, while $\omega_3$ was poorly recovered, as reported in previous work (Reed et al., 2020; Hein et al., 2021). See figure below:

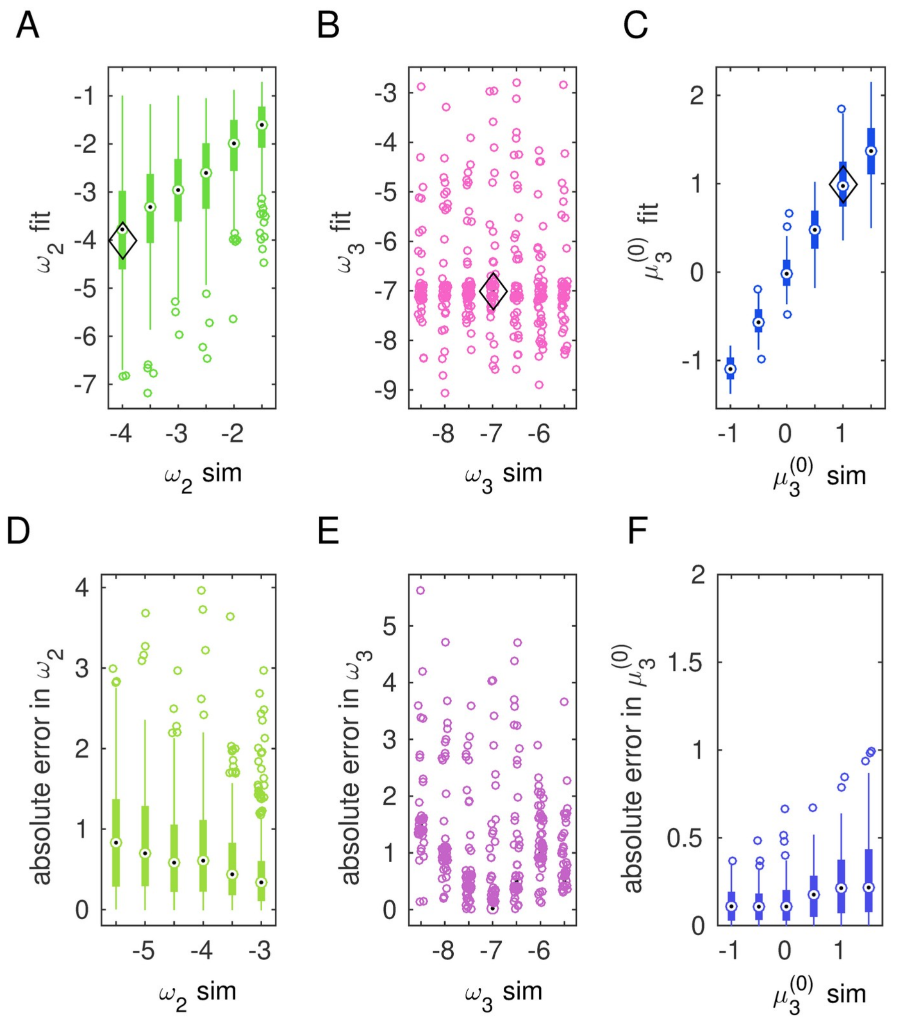


**HGF parameter estimation using the input observed by one participant (ID zhanx_25). A-C)** Boxplots (median, 25 and 75 percentiles) illustrating the results of parameter estimation for $\omega_2$ (A), $\omega_3$ (B) and $\mu_3^{(0)}$ (C). The x-axis represents the set parameters introduced in the simulated responses (labelled “sim”), while y-axis data reveal the corresponding estimated value of that same parameter (labelled “fit”). Parameters $\omega_2$ and $\mu_3^{(0)}$ were estimated with high accuracy, as there was a high significant correlation between simulated and estimated (fit) values: Pearson R = 0.6306, $P << 1 \times 10^{-6}$ for $\omega_2$, R = 0.9656, $P << 1 \times 10^{-6}$ for $\mu_3^{(0)}$. Parameter $\omega_3$ was poorly estimated: R = -0.0242, $P$ = 0.2668. The prior values of $\omega_2$, $\omega_3$ and $\mu_3^{(0)}$ used in the configuration file for estimating each parameter from the simulated responses were as defined in **Table S1** for the best fitting model (HGF$_{\mu 3}$): $\omega_2$ = -4, $\omega_3$ = -7, respectively (variance 16 in both cases); $\mu_3^{(0)}$ = 1 (variance 1). Prior values are denoted by the diamond shape in the top panels.

A complementary analysis using simulated responses to observed inputs from a different participant (ID zhanx_3) provided similar results.

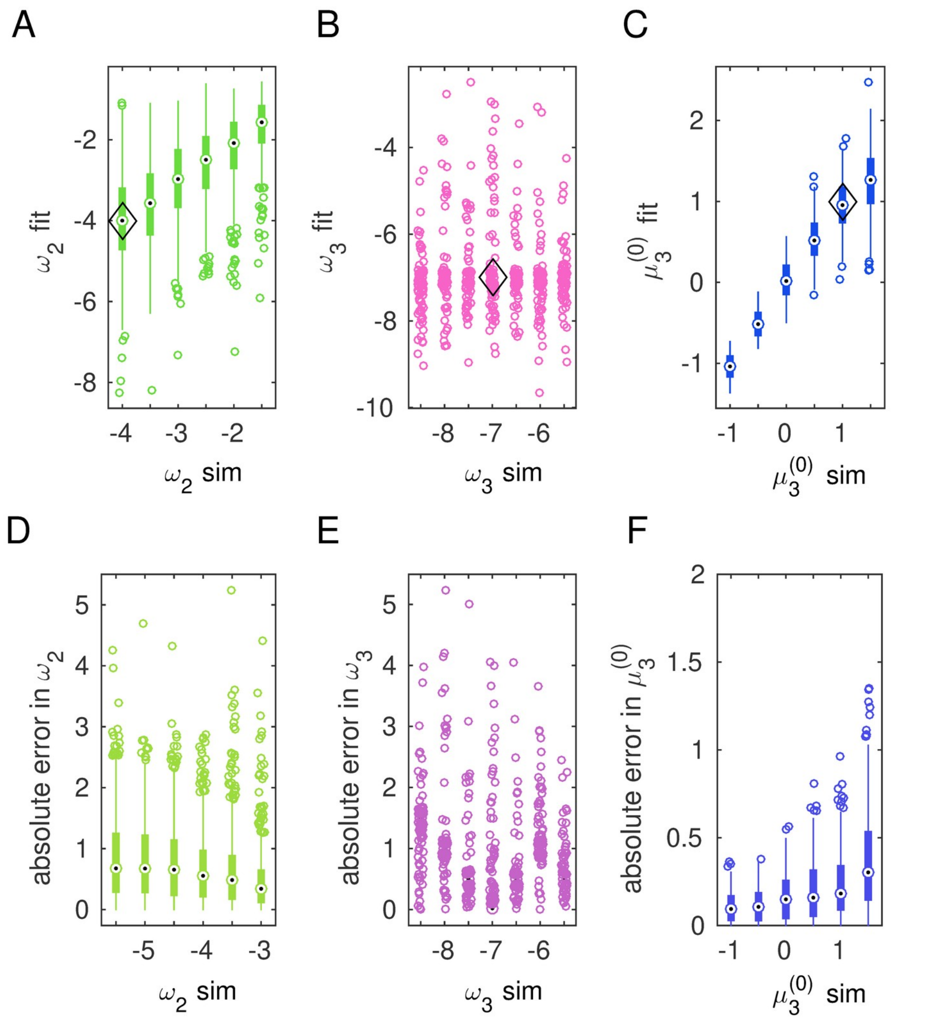


**HGF parameter estimation using the input observed by one participant (ID zhanx_3).** Same as the figure above. Estimation of parameters $\omega_2$ and $\mu_3^{(0)}$ was highly accurate: Correlation between simulated and fitted value Pearson R = 0.6512, $P \ll 1 \times 10^{-6}$ for $\omega_2$, R = 0.9468, $P \ll 1 \times 10^{-6}$ for $\mu_3^{(0)}$. Parameter $\omega_3$ was also poorly estimated: R = 0.0019, $P$ = 0.9019.

### Table S1

| Model | Prior | Mean | Variance |
|---|---|---|---|
| **$HGF_3$** | κ | 1 | 0 |
| | $\omega_2$ | −4 | 16 |
| | $\omega_3$ | −7 | 16 |
| | $\mu_2^{(0)}$ | 0 | 0 |
| | $\sigma_2^{(0)}$ | 0.1 | 0 |
| | $\mu_3^{(0)}$ | 1 | 0 |
| | $\sigma_3^{(0)}$ | 1 | 0 |
| | ζ | 48 | 1 |
| **$HGF_2$** | κ | 0 | 0 |
| | $\omega_2$ | −4 | 16 |
| | $\omega_3$ | −7 | 0 |
| | $\mu_2^{(0)}$ | 0 | 0 |
| | $\sigma_2^{(0)}$ | 0.1 | 0 |
| | $\mu_3^{(0)}$ | 1 | 0 |
| | $\sigma_3^{(0)}$ | 1 | 0 |
| | ζ | 48 | 1 |
| **$HGF\mu_3$** | κ | 1 | 0 |
| | $\omega_2$ | −4 | 16 |
| | $\omega_3$ | −7 | 16 |
| | $\mu_2^{(0)}$ | 0 | 0 |
| | $\sigma_2^{(0)}$ | 0.1 | 0 |
| | $\mu_3^{(0)}$ | 1 | 1 |
| | $\sigma_3^{(0)}$ | 1 | 1 |

**Table S1.** Means and variances of the priors on perceptual parameters and starting values of the beliefs of the HGF models. Values are shown for 3-level HGF, 2-level HGF and $HGF_{\mu3}$ models. Free parameters are estimated in their unbounded space. Accordingly, parameters that are restricted to a confined interval are log-transformed, to allow for estimation in an unbounded space. As in recent work (Weber et al., 2020; Hein et al., 2021), the initial values of the belief trajectories were fixed in each individual for the 3-level HGF and 2-level HGF models: $\mu_2^{(0)}$, $\sigma_2^{(0)}$, $\mu_3^{(0)}$, $\sigma_3^{(0)}$. We estimated $\omega_2$, $\omega_3$ in each participant ($\omega_2$ only for the 2-level HGF). In addition, parameter ζ was estimated in the log space (3-level HGF and 2-level HGF models). The winning model $HGF_{\mu3}$ had as free parameters $\omega_2$, $\omega_3$ , $\mu_3^{(0)}$, and $\sigma_3^{(0)}$ , and the mapping from beliefs to decisions was a function of the inverse decision noise paremeter $e^{-\mu_3^{k-1}}$ (Diaconescu et al., 2014). Here, $\sigma_3^{(0)}$ was estimated in the log space.
